# Discovery of a novel antibiotic class targeting the enolase of *Acinetobacter baumannii*

**DOI:** 10.1101/2025.04.23.650324

**Authors:** Irene Molina Panadero, Antonio Moreno Rodríguez, Mercedes de la Cruz, Pilar Sánchez, Laura Tomás Gallardo, Thanadon Samernate, Milan Sencanski, Sanja Glisic, Olga Genilloud, Poochit Nonejuie, Antonio J. Pérez-Pulido, Abdelkrim Hmadcha, Younes Smani

**Affiliations:** Centro Andaluz de Biología del Desarrollo, Universidad Pablo de Olavide/CSIC/Junta de Andalucía, 41013 Seville, Spain; Fundación MEDINA, Parque Tecnológico Ciencias de la Salud, 18016 Granada, España; Proteomics and Biochemistry Platform. Centro Andaluz de Biología del Desarrollo. Universidad Pablo de Olavide/CSIC/Junta de Andalucía, 41013 Seville, Spain; Institute of Molecular Biosciences, Mahidol University, Salaya, Nakhon Pathom 73170, Thailand; Laboratory for Bioinformatics and Computational Chemistry, Institute of Nuclear Sciences VINCA, National Institute of the Republic of Serbia, University of Belgrade, 11001 Belgrade, Serbia; Departamento de Biología Molecular e Ingeniería Bioquímica, Universidad Pablo de Olavide, 41013 Seville, Spain; Biosanitary Research Institute (IIB-VIU), Valencian International University (VIU), 46021 Valencia, Spain

**Keywords:** ENOblock, enolase, synergy, high-throughput screening, drug repurposing, bacteria, infection, *Acinetobacter baumannii*

## Abstract

High-throughput screening studies provide an additional approach to discovering repurposed drugs for antimicrobial treatments. In this work, we report the identification of ENOblock, an anticancer drug, as a novel antibiotic class. We computationally and experimentally validated that ENOblock synergizes with colistin, the last resort antibiotic, the colistin. Additionally, we identified enolase as the potential bacterial target for ENOblock. The *in silico* and *in vitro* antibacterial activity of ENOblock translated into potent *in vivo* efficacy in animal infection model. Collectively, the preclinical data support the selection of ENOblock as a promising candidate for antimicrobial development, with the potential to address the urgent threat of infections caused by *Acinetobacter baumannii*.

## INTRODUCTION

Gram-negative bacteria (GNB) are a significant concern in healthcare settings due to their ability to cause a wide range of infections such as pneumonia, bloodstream infections, wound or surgical site infections, and meningitis^1^, which are often difficult to treat. GNB infections, especially hospital acquired ones, represent a significant burden on healthcare systems worldwide, with reported costs of $136 million per year^2^, requiring ongoing efforts in surveillance, infection prevention and control, antibiotic stewardship, and research into new treatment options to mitigate their impact.

One of the main difficulties in tackling GNB is their high efficiency in acquiring antimicrobial resistance (AMR) encoded by genomic, transcriptomic and proteomic changes^3^. Compounding the problem of AMR with reported high number of deaths associated with bacterial AMR and/or attributable to bacterial AMR is the immediate threat of a reduction in the discovery and development of new antibiotics. The World Health Organization (WHO)^4^ and other European institutions have recently underscored the dangers posed by these infections^5^. Consequently, a perfect storm is converging regarding these infections: increasing antimicrobial resistance with a decreased new drug development^5^. This context is likely the best example of the purported “Post-Antibiotic Era”, with relevance even in non-specialized media. It is clear that new policies and actions are necessary to avoid the forecasts for 2050 that attribute ten million deaths worldwide to antimicrobial resistance^5^. Especially for pathogens like *Acinetobacter baumannii* that is reported with continuous increase in AMR and are accounted for most reported associated diseases and deaths^6^.

Approaches using computational methods and high-throughput screening (HTS) have recently been developed for antibiotic discovery^7–10^. For example, screening small-molecule libraries has revealed new antimicrobial agents that belong to existing or new antibiotic classes^11–13^. Recently, HTS studies have been developed to discover repurposed drugs for antimicrobial treatments^14^.

Specific enolase inhibitors, such as 2-aminothiazoles, directly impair its catalytic activity, disrupting ATP production and compromising bacterial viability^15^. The evolutionary conservation of this enzyme in multiple Gram-negative pathogens underscoress its potential as a therapeutic target, while its multifunctional role in virulence further enhances its clinical relevance^16^.

In this work, we report the identification of ENOblock, an anticancer drug, as a novel antibiotic class. We computationally and experimentally validated that ENOblock synergizes with colistin the last resort antibiotic, the colistin. Additionally, we identified enolase as the potential bacterial target for ENOblock. The *in silico* and *in vitro* antibacterial activity of ENOblock translated into potent *in vivo* efficacy in the *Galleria mellonella* animal infection model. Collectively, these preclinical data could support the selection of ENOblock as a promising candidate for antimicrobial development with the potential to address the urgent threat of infections caused by *A. baumannii*.

## MATERIAL AND METHODS

### Bacterial Strains

A total of 32 clinical strains of *A. baumannii* colistin-resistant (n=14)^17,18^ or carbapenem-intermediate/resistant (n=18) were collected from the “II Spanish Study of *A. baumannii* GEIH-REIPI 2000-2010” multicenter study (GenBank Bioproject PRJNA422585) and the reference strain Ab ATCC 17978 were used in this study.

HTS was performed with two reference strains of *A. baumannii* and *Escherichia coli* (Ab ATCC 17978 and ATCC 25922, respectively) and five clonally unrelated clinical strains:

*A. baumannii* Ab9 (ST672), colistin- and tigecycline-susceptible; MDR *A. baumannii* Ab186 (ST208), colistin-susceptible, tigecycline-resistant; *E. coli* C1-7-LE (ST8671), colistin- and tigecycline-susceptible; and MDR *E. coli* MCR1^+^ (ST6108), colistin-resistant, tigecycline-susceptible^19^; *Klebsiella pneumoniae* Kp10 KPC-2 producing^20^

### EU-OPENSCREEN library and HTS validation

EU-OPENSCREEN provided a subset of 2,464 bioactive compounds from the ECBL pilot library (<u>EU-OPENSCREEN: European Chemical Biology Library - Pilot Library</u>) as 10 mM stock solutions in 100% DMSO. The library was screened in duplicate at a final concentration of 100 μM per well (1% DMSO). The antibacterial single-concentration HTS and dose-response susceptibility assays were conducted in 384-well plates, with bacterial cell density measured by optical density at 600 nm (OD600).

A starting inoculum of 10^6^ colony-forming units (CFU)/mL was used, with an incubation time of 24 hours for the *A. baumannii* (Ab ATCC 17978, Ab9, and Ab186) strains and *E. coli* (ATCC 25922, C17LE, and MCR1^+^) strains. Imipenem and colistin were used as internal controls for *A. baumannii* and *E. coli* strains, respectively.

Compounds were distributed in duplicate on separate 384-well microtiter plates using an Echo 550® acoustic liquid handler (Beckman Coulter™, Indianapolis, IN) and inoculated with bacteria to a final concentration of 10^6^ CFU/mL, with a total assay volume of 25.25 μL. Plates were incubated with shaking for 24 hours at 37 °C. Bacterial growth was measured by reading the OD600 using an EnVision™ microplate reader (Revvity, Waltham, MA). The activity of the compounds was expressed as the percentage of bacterial growth inhibition, and it was calculated using the following normalization:

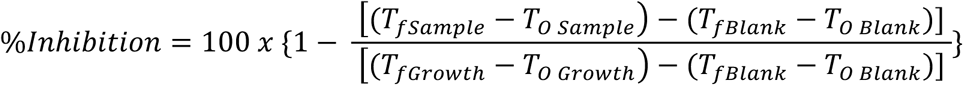

Where, **T0 Sample** is the absorbance of the strain growth in the presence of compound measured at time zero, **Tf Sample** is the absorbance of the strain growth in the presence of compound measured at final time,**T0 Growth** is the absorbance of the strain growth in the absence of compound measured at time zero, **Tf Growth** is the absorbance of the strain growth in the absence of compound measured at final time, **T0 Blank** is the absorbance of the broth medium (blank) measured at time zero, **Tf Blank** : the absorbance of the broth medium (blank) measured at final time**; T0** is Time at 0 hour and **Tf** is Time at 24 hours. **Blank:** is composed of 25 μL of MHII and 0.25 μL of DMSO 20%

**Growth:** is composed of 25 μL of bacterial inoculum and 0.25 μL of DMSO 20%.

**Sample:** studied compound.

The Genedata Screener software (Genedata, Inc., Basel, Switzerland) was used to process and analyse all the screening data. Their reproducibility and sensitivity were supported by the statistical values derived from all the experiments performed. Also, MIC 90% (Minimum Inhibitory Concentration required to inhibit 90% of the growth of a microorganism) value for every Dose Response Curve of reference antibiotic compounds (imipenem and colistin) was determined to assess consistent reproducible activity data within assay plates and between experiments. This software was used to calculate quality control parameters such as RZ’ factor (RZ’ factor ≥0.5), and signal/background ratio (S/B). The Z’factor predicts the robustness of an assay by considering the mean and standard deviation of both positive and negative controls^21^. The robust Z’ factor (RZ’ factor) is based on the Z’ factor, but standard deviations and means are replaced by the robust standard deviations and medians, respectively.

### *in vitro* susceptibility testing

The minimum inhibitory concentration (MIC) of ENOblock and colistin were determined against all studied *A. baumannii*, *E. coli* and *Klebsiella pneumoniae* strains in two independent experiments using the broth microdilution method, following the standard guidelines of the European Committee on Antimicrobial Susceptibility Testing (EUCAST)^22^. A 5×10^5^ CFU/mL inoculum of each strain was cultured in Luria-Bertani (LB) and cation-adjusted Mueller-Hinton broth, and then added to U-bottom microtiter plates (Deltalab, Spain) containing ENOblock or colistin. The plates were incubated for 18 hours at 37°C. *Pseudomonas aeruginosa* ATCC 27853 was used as the positive control strain.

### Antimicrobial selection pressure

The diluted Ab ATCC 17978 inocula (10^5^ CFU/mL) were incubated with sub-inhibitory concentrations of ENOblock, corresponding to dilutions one-fold below the MIC, at 37°C for 24 hours. Bacterial concentration was determined in cultures showing positive growth, which were then re-adjusted to 10^5^ CFU/mL for further incubation with a two-fold increased concentration of ENOblock. These steps were repeated until an ENOblock concentration was reached that completely inhibited bacterial growth.

### Time kill kinetic assays

To determine the bactericidal activity, duplicate time-kill curves were performed for *A. baumannii* ATCC 179178 and Ab CR17 strains, for *E. coli* MCR1^+^ (colistin-resistant) and *K. pneumoniae* Kp10 (carbapenem-resistant) strains, and for *E. coli* MG1655 and its isogenic deficient in enolase (Ec Δ*eno*) strain^23^, as previously described^24^. An initial inoculum of 5×10^5^ CFU/mL was added to LB in the presence of 1xMIC, 2xMIC and 4xMIC of ENOblock. A drug-free broth was evaluated in parallel as a control. Tubes of each condition were incubated at 37 °C with shaking, and viable counts were determined by serial dilution at 0, 2, 4, 8, and 24 hours. Viable counts were determined by plating 100 µL of the control, test cultures, or the respective dilutions at the indicated times onto sheep blood agar plates (ThermoFisher, Spain). Plates were incubated for 24 hours at 37 °C, and after colony counts, the log10 of viable cells (CFU/mL) was determined. Bactericidal activity was defined as a reduction of ≥3 log10 CFU/mL from the initial inoculum.

### EIIP/AQVN filter

Specific recognition and targeting between interacting biological molecules at distances > 5 Å were determined by the average quasi-valence number (AQVN) and the Electron-ion interaction potential (EIIP) derived from the general model pseudopotential^25^

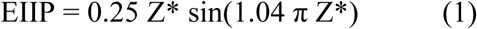

where Z* is the AQVN determined by:

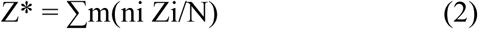

where Zi is the valence number of the ith atomic component, ni is the number of atoms of the ith component, m is the number of atomic components in the molecule, and N is the total number of atoms. EIIP values are computed using Equations (1) and (2) and are expressed in Rydberg units (Ry).

AQVN and EIIP are unique physical properties that characterize long-range interactions between biological molecules among the 3300 molecular descriptors currently in use^26^. It has been shown that the EIIP and AQVN of organic molecules strongly correlate with their biological activity (mutagenicity, carcinogenicity, toxicity, antibiotic and cytostatic activity, etc.)^27^.

### Checkerboard assay

The assay was performed on a 96-well plate in duplicate as previously described^24^. Colistin, imipenem, ceftazidime or tigecycline were two-fold serially diluted along the x-axis, whereas ENOblock was two-fold serially diluted along the y axis to create a matrix, where each well consists of a combination of both agents at different concentrations. Bacterial cultures grown overnight were then diluted in saline to 0.5 McFarland turbidity, followed by 1:50 further dilution LB and inoculation on each well to achieve a final concentration of approximately 5.5×10^5^ CFU/mL. The 96-well plates were then incubated at 37°C for 18 hours and examined for visible turbidity. The fractional inhibitory concentration (FIC) of the colistin, imipenem, ceftazidime or tigecycline was calculated by dividing the MIC of colistin, imipenem, ceftazidime or tigecycline in the presence of ENOblock by the MIC of colistin, imipenem, ceftazidime or tigecycline alone. Similarly, the FIC of ENOblock was calculated by dividing the MIC of ENOblock in the presence of imipenem, ceftazidime or tigecycline alone. The FIC index was the summation of both FIC values. FIC index values of ≤0.5 and >5 were interpreted as synergistic and non-synergistic, respectively.

### Bacterial growth curves

To confirm the synergy between ENOblock and colistin, bacterial growth curves of the *A. baumannii* ATCC 17978 and Ab CR17 strains were performed in duplicate in 96-well plate (Deltlab, Spain). An initial inoculum of 5×10^5^ CFU/mL was prepared in LB in the presence of colistin (0.12 or 1 mg/L) and ENOblock (8 mg/L) separately or together. A drug-free broth was evaluated in parallel as a control. Plates were incubated at 37°C with shaking, and bacterial growth was monitored for 24 hours using a microtiter plate reader (Tecan Spark, Austria).

### Bacterial cytological profiling

#### Fluorescent microscopy

Overnight cultures of *A. baumannii* ATCC 17978 were diluted 1:100 in LB broth and incubated on a roller at 30°C until the OD600 reached 0.2. ENOblock was added to the bacterial cultures at MIC levels prior to observation under a fluorescence microscope at various time points. After each timepoint, the cultures were stained with 2 µg/mL FM4-64, 2 µg/mL DAPI, and 0.5 µM SYTOX Green. The bacterial cells were then harvested by centrifugation at 6,000 x *g* for one minute and resuspended in 1/10 of the original volume. A small amount of the concentrated bacterial cultures was placed on an agarose pad (1.2% agarose in 10% LB broth) on concave glass slides for microscopy. Consistent experimental settings and imaging parameters were maintained throughout all experiments included in the statistical analysis of the antibiotic training sets.

#### Image data analysis

The raw images from the fluorescent microscope were preprocessed using ImageJ software^28^, and cell features were extracted with CellProfiler 4.0 software^29^. After data extraction, the ENOblock-treated cell profiles were analyzed alongside antibiotic-treated cell profiles from previous studies using an analysis pipeline from previous research^30^. Briefly, the data was transformed using QuantileTransformer^31^, and outliers were removed using hierarchical density-based spatial clustering of applications with noise (HDBSCAN)^32^. This analysis utilized a morphological feature set from previous studies^30^. Finally, the dimension of the data set was reduced and visualized through data clustering using pairwise controlled manifold approximation (PaCMAP)^33^.

### Membrane permeability assay

Membrane permeability of *E. coli* MCR1^+^ and *K. pneumoniae* Kp10 strains was assessed in the presence of ENOblock following a previously established protocol^24^. Briefly, bacterial cells were grown in LB broth overnight and bacterial pellets were harvested by centrifugation at 4,600 *g* for 15 minutes, washed with 1X PBS, and centrifuged again under the same conditions. The bacterial pellets OD were adjusted to 0.2 and incubated with 0.5xMIC ENOblock. Then, they were resuspended in 100 µL of 1X PBS containing 10 μL of Ethidium Homodimer-1 (EthD-1) (ThermoFisher, Spain). After 10 minutes of incubation, 100 µL of the suspension was transferred to a 96-well plate, and fluorescence was measured over 3 hours using a Typhoon FLA 9000 laser scanner (GE Healthcare Life Sciences, USA). Fluorescence intensity was quantified using ImageQuant TL software (GE Healthcare Life Sciences, USA).

### Docking and molecular modelling

The crystal structure of enolase C-terminal of *A. baumannii* was predicted by AlphaFold^34^. This target protein was modified using Autodock tools 1.5.6 software; including ligand and water removal, hydrogen addition, and incorporation of Kollman charges. The resultant files were saved in pdbqt format. The ligand “ENOblock” was downloaded from Pubchem (https://pubchem.ncbi.nlm.nih.gov/) in SMILES format (https://pubchem.ncbi.nlm.nih.gov/compound/24012277). Openbabel 3.1.1 software was used to create the 3D chemical structure, energy minimization, hydrogen atoms addition and establishment of a neutral pH. Resulting ligand was saved in MOL.2 format. Gasteiger charges computation for the ligand structure was performed using Autodock tools 1.5.6, with the output saved in pdbqt format. Autodock tools 1.5.6 was used for docking and subsequent analysis of the docking results.

### Protein expression and purification of recombinant AbEnolase

The *A. baumannii* ATCC 17978 enolase wild-type gene (UniProtKB: A3M5Y1), enolase mutant #1 (D207A: aspartate replaced by alanine at position 207), enolase mutant #2 (S371A: serine replaced by alanine at position 371) and enolase double mutant (D207A and S371) were synthesized and cloned into the pET-30a(+) vector with an N-terminal His6 tag using *NdeI* and *XhoI* restriction sites by GenScript. Prior to synthesis, the codon usage of the gene was optimized using the GenSmart Tool. *Wt* and mutated AbEnolase overexpression were performed in *E. coli* Rosetta™ DE3 pRare (Merck Millipore, Spain). A single colony from freshly transformed cells was incubated overnight at 37°C in LB supplemented with ampicillin, with shaking at 180 rpm. The culture was diluted 1:100 into fresh medium and grown at 37°C until an OD600 of 0.5 was reached. Protein expression was induced with 0.5 mM isopropyl-β-D-thiogalactopyranoside (IPTG) (Merck, Spain) for 3 hours at 30°C. Cells were harvested by centrifugation, washed with 50 mM TrisHCl pH 8 and the pellet was stored at -80°C until use. The cell pellet was resuspended in Lysis Buffer (50 mM Tris, pH 8.0, 0.4 M NaCl, 5 mM imidazole, 5 mM MgCl2) and disrupted by sonication (Branson SFX550 sonifier, Emerson, Spain). Cell debris was removed by centrifugation at 10,000 *g* for 25 minutes. The soluble lysate was filtered through a 0.45 μm filter and applied to a nickel affinity column HisTrap FF (Cytiva, USA) equilibrated with Lysis Buffer using an ÄKTApure™ chromatography system (Cytiva, USA). Enolase was eluted using a 10-column volume gradient of lysis buffer supplemented with 250 mM imidazole. Fractions containing enolase were identified by SDS-PAGE, pooled, and dialyzed overnight at 4°C against Dialysis Buffer (25 mM Tris, pH 7.5, 100 mM NaCl, 5 mM MgCl2, 0.5 mM dithiothreitol [DTT]) using a 3.5K Slide-A-Lyzer Dialysis Cassette (Thermo Scientific, Spain). Before storage at -80°C, the protein was concentrated using a 10K Amicon filter (Millipore, Spain) and quantified by densitometric analysis. Aliquots were prepared at a concentration of 5 mg/mL and stored at -80°C. See **Figure S1**.

### Isothermal titration calorimetry (ITC)

Interaction of the wt Abenolase and mutants D207A, S371A and D207A+S371S with ENOblock was analyzed with an Auto-iTC200 isothermal titration calorimeter (MicroCal, Malvern-Panalytical) at a constant temperature of 25°C. Protein solution was used at a final concentration of 75 μM in buffer 25 mM Tris pH 7.5, 100 mM NaCl, 5 mM MgCl2. ENOblock solution at 1 mM in the same buffer was injected following programmed sequences of 13 injections of 3 μL with a spacing of 150 s and a stirring speed of 750. Three independent calorimetric titrations (n = 3) were performed for each interaction, in order to assess reproducibility, as well as to estimate average values for the dissociation constants and their errors. The dissociation constants were obtained through nonlinear least squares regression analysis of the experimental data to a model considering a single ligand binding site. Appropriate controls (buffer into buffer; ligand dilution into buffer; buffer into enolase) were conducted to obtain a baseline for each experiment and determine the heat of dilution/mixing.

### Generation of enolase knockout from *A. baumannii* Ab ATCC 17978

To construct an *enolase* knockout from *A. baumannii* Ab ATCC 17978, we followed the protocol described previously^35,36^. Briefly, an internal *enolase* 482-bp fragment obtained by PCR amplification with the primers enolase IntUp and enolase IntLw (**Table S1**) was cloned into pGEM-T (Promega, Spain) to give plasmid enolase-pGEM-T by using T4 DNA ligase (Promega, Spain). The resulting construct incorporated into *E. coli* DH5α was purified and electroporated into Ab ATCC 17978 to knock out the *enolase* gene.

Transformants were selected on LB agar plates containing 80 µg/mL ticarcillin. The *enolase* gene disruption within the resulting strain, designated Ab Δ*eno*, was confirmed by PCR using a combination of primers matching the upstream region of *enolase* gene and the pGEM-T Easy vector.

### Bacterial enolase activity assay

Enolase activity was determined in three independent experiments following the instructions of Enolase Activity Assay kit (Merck, Spain). Briefly, a bacterial inoculum of 10^6^ CFU/mL, after centrifugation, was incubated with a master mix composed by enolase substrate, peroxidase substrate, and the necessary converters for the colorimetric reaction at 25°C. Enolase enzymatic activity, reported in mU/mL, was measured at OD570nm every two-three minutes for one hour using a microtiter plate reader (Tecan Spark, Austria) and calculated using the specific formula: ΔA570 = (A570)final - (A570)initial, and the following equation: enolase Activity = (B × Sample Dilution Factor) / (Reaction Time × V) where: **B** = Amount (nmol) of H2O2 generated between Tinitial and Tfinal, **Reaction Time** = Tfinal – Tinitial (minutes), and **V** = volume of sample (mL) added to the well

### Bacterial growth curves

To determine the antibacterial of ENOblock against *wild-type* and enolase-deficient *A. baumannii*, bacterial growth curves of the *A. baumannii* Ab ATCC 17978 and its isogenic deficient in enolase (Ab Δ*eno*) strain in similar conditions (see above). Bacteria was incubated in the presence of 1x, 2x and 4x MIC of ENOblock.

### Human cell culture

HeLa cells were grown in DMEM supplemented with 10% heat-inactivated fetal bovine serum (FBS), vancomycin (50 mg/L), gentamicin (20 mg/L), and amphotericin B (0.25 mg/L) (Invitrogen, Spain), and 1% HEPES in a humidified incubator with 5% CO2 at 37°C. The HeLa cells were routinely passaged every 3 or 4 days. Immediately before infection, HeLa cells were washed three times with prewarmed PBS and further incubated in DMEM without FBS and antibiotics^37^. The human monocytic leukemia cell line THP-1 (ATCC, LGC Standards, Spain) was cultured in RPMI medium (ThermoFisher, Spain) supplemented with 10% FBS (ThermoFisher, Spain), 1% penicillin-streptomycin (Gibco, Spain), 1% HEPES, and 0.05 mM 2-mercaptoethanol (Gibco, Spain). Cells were maintained in a humidified incubator at 37°C with 5% CO₂ and were routinely passaged every 3 days.

### Differentiation of THP-1 monocytes into macrophages

To differentiate THP-1 monocytes into macrophages, the cells were seeded in RPMI medium supplemented with 10% FBS, 1% penicillin-streptomycin, and 40 ng/mL phorbol 12-myristate 13-acetate (PMA) (Merck, Spain). The cells were incubated at 37°C with 5% CO₂ for 2 days^38^. Before infection, macrophages were washed three times with prewarmed PBS and further incubated in RPMI medium without FBS or antibiotics^37^.

### Tandem Mass Tag (TMT) assay and analysis

Infected and non-infected HeLa and macrophage cells were lysed in a buffer containing 4% SDS, 100 mM Tris (pH 7.6), and water. Cell membranes were disrupted via sonication for 1.5 minutes with 5-second intervals on ice. The supernatant was collected after centrifugation at 4°C for 5 minutes. Protein concentration was determined using a DC Protein Assay Kit (Bio-Rad, Spain). Samples were incubated with 10 mM TCEP in 100 mM Tris (pH7.6) at 55°C for 1 hour and then treated in darkness with 375 mM iodoacetamide in 100 mM TrisHCl (pH7.6) for 30 minutes. Proteins were precipitated with acetone and stored at -20°C for at least 4 hours. Samples were centrifuged at 16.000 *g* at 4°C for 10 minutes, and the acetone was discarded by pellet desiccation. To analyse differential protein expression between bacteria alone and bacteria in contact with HeLa or macrophage cells, an isobaric standard tandem tag (ThermoFisher, Spain) was employed. Samples were analyzed via tandem mass spectrometry (MS/MS) (Bruker, USA) at the Proteomics Facility of the University Pablo de Olavide (Seville, Spain). Protein pellets were resuspended in triethylammonium bicarbonate and digested overnight at 37 °C using trypsin bovine (Sequencing Grade Modified Trypsin, Promega) in a ratio 1:12 enzime-substrate. Reaction was stopped using formic acid to 0.5% and samples were labelled with the isobaric tags following manufacturer instructions, using channels 126, 127, 128, 129, 130 and 131. 5ug of every tagged sample were mixed in a single sample tube. OMIX C18 tips (Agilent Technologies) were used for concentrating and desalting tagged peptide extracts. Sample was dried and resuspended in 0.1% trifluoroacetic acid and injected in nano-HPLC system. Protein digested samples were separated in a Thermo ScientificTM Easy nLC system using a 50cm C18 Thermo Scientific™ EASY-Spray™ column. The following solvents were employed as mobile phases: Water 0.1% Formic Acid (phase A) and Acetonitrile, 20% H2O, 0.1% Formic Acid (phase B). Separation was achieved with an acetonitrile gradient from 10% to 35% over 360 min, 35% to 100% over 1 min, and 100% B over 5 min at a flow rate of 200 nL/min.

A Thermo ScientificTM Q Exactive™ Plus Orbitrap™ mass spectrometer was used for acquiring the top 10 MS/MS spectra in DDA mode. LC-MS data were analysed using the SEQUEST® HT search engine in Thermo Scientific™ Proteome Discoverer™ 2.2 software considering the modifications: static carbamidomethylation (C), dynamic oxidation (M) and dynamic N-terminus acetylation. Data were searched against the Uniprot *Acinetobacter_baumanii* or *Homo sapiens* (sp_canonical TaxID=9606) (v2024-03-27) protein databases and results were filtered using a 1% protein FDR threshold. Quantifiable proteins were those identified by >2 peptides with a confidence level >95%, a p-value <0.05, and an error factor <2 for each reference tag. Proteins with a fold change of <-1 or >1 were classified as downexpressed or overexpressed, respectively. Sample-to-sample distances were visualized through principal component analysis (PCA). Functional enrichment analyses included Clusters of Orthologous Groups (COG) and Gene Onthology (GO) terms that were assigned using EggNOG mapper. Enriched GO terms were identified using the enricher function in the topGO R package.

### Adhesion and invasion assays

HeLa and macrophage cells were infected with *A. baumannii* Ab ATCC 17978 and Ab CR17 strains, and for *E. coli* MCR1^+^ and *K. pneumoniae* Kp10 strains at a concentration of 1×10^8^ CFU/mL, in the absence and presence of 1xMIC of ENOblock at a multiplicity of infection (MOI) of 100. The infection was carried out for two hours with 5% CO2 at 37°C in three independent experiments. After that, the infected HeLa and macrophage cells were washed five times with pre-warmed PBS and lysed with 0.5% Triton X-100. Diluted lysates were plated onto LB agar and incubated at 37°C for 24 hours to enumerate the developed colonies and determine the number of bacteria that had attached to the HeLa and macrophage cells^37^. In addition, to determine the number of colonies that entered inside the HeLa and macrophage cells, the wells were washed with phosphate buffered saline and incubated for 30 minutes in the presence of DMEM or RPMI plus gentamicin (256 µg/mL), in order to kill the bacteria, present in the area. Then, the wells were washed with phosphate buffered saline to remove gentamicin. The number of colonies that entered inside HeLa and macrophage cells was determined as described above^37^.

### Cellular toxicity of ENOblock

HeLa cells and macrophages differentiated from THP-1 cells^38^ were incubated with ENOblock at different concentrations ranged from 0.5 to 256 mg/L for 24 hours with 5% CO2 at 37°C. Prior the evaluation of the ENOblock cytotoxicity, HeLa and macrophage cells were washed three times with prewarmed PBS 1X. Subsequently, quantitative cytotoxicity was evaluated by measuring the mitochondrial reduction activity using the 3-(4,5-dimethylthiazol-2-yl)-2,5-diphenyltetrazolium bromide (MTT) assay as described previously^35^. The percentage of cytotoxicity was calculated from the absorbance at 570 nm as follow: [(Absorbance 570 nm of treated cells/Absorbance 570 nm mean of untreated cells) × 100]. The cytotoxic concentration 50% (CC50) value was determined using GraphPad Prism 9.

### Immunofluorescence

Immunofluorescence assay was performed as described previously^39^. Briefly, the HeLa cells plated on coverslips were incubated with *A. baumannii* CR17 and *E. coli* MCR1^+^ strains for two hours, and later were incubated with ENOblock (0 and 1xMIC, 30 minutes) at 5% CO2 and 37°C. Bacterial cells were removed, and HeLa cells were washed five times with cold PBS. HeLa cells on the coverslips were fixed in methanol for 8 minutes at -20 °C, permeabilized with 0.5% Triton X-100 and blocked with 20% pork serum in PBS. Primary antibodies: anti-OmpA of *A. baumannii* (ThermoFischer, Spain), mouse anti-*E. coli* (Abcam, Spain), and rabbit anti-human fibronectin (Merck, Spain) were used at dilution of 1:25, 1:25 and 1:50, respectively, in PBS containing 1% bovine serum albumin (BSA) for 2 hours. After washing with PBS, the coverslips were incubated with their respective secondary antibodies: Alexa488-conjugated goat anti-mouse IgG, and Alexa594-conjugated goat anti-rabbit IgG (Invitrogen, Spain) at dilution of 1:50, 1:50 and 1:100, respectively, in PBS containing 1% BSA for 1 hour. The fixed coverslips were incubated for 10 minutes at room temperature with DAPI (Applichem, Germany) (0.5 μg/mL), washed with PBS, mounted in fluorescence mounting medium “Prolong Diamond Antifade Mounting” (Invitrogen, Spain), and visualized using fluorescence microscopy Zeiss Axio Imager 2 (Zeiss, Germany).

### Informational Spectrum Method

The Informational Spectrum Method (ISM), a virtual spectroscopy approach, was employed to investigate protein-protein interactions *in silico* to complement experimental data. ISM converts protein or DNA sequences into numerical signals derived from the EIIP of their constituent amino acids or nucleotides. EIIP values capture electronic properties that mediate long-range molecular interactions (5 to 1000 Å)^40^. This virtual spectroscopy method enables functional analysis of protein sequences without requiring prior experimental input. The ISM extension for small molecules (ISM-SM) uses similar principles, translating small molecules represented in SMILES notation into EIIP-based arrays corresponding to atomic groups^41^. By identifying shared frequencies in computational spectra, ISM-SM predicts potential interactions between proteins and small molecules and highlights likely binding regions on the protein. In this study, ISM-SM was applied to identify shared informational features among *A. baumannii* enolase (UniProt: sp|B0VQI4|ENO_ACIBS), human plasminogen (UniProt: sp|P00747|PLMN_HUMAN), human fibrinogen (UniProt: sp|P02679|FIBG_HUMAN), human fibronectin (UniProt: sp|P02751|FINC_HUMAN) and ENOblock (PubChem CID: 24012277). In the next step, the plasminogen-enolase, fibrinogen-enolase, and fibronectin-enolase complexes were modelled. The obtained sequences of enolase and these extracellular matrix (ECM) proteins were sent to Alfafold3 to build a protein-protein complex.

### Coating human host proteins on wells

The coating of human host proteins (fibrinogen, fibronectin, and plasminogen) onto 96-well plates was performed as previously described^42^. Wells were coated overnight at 4°C with 125 µL of PBS containing 1.25 µg of plasmatic plasminogen, fibronectin or fibrinogen (10 µg/mL), or with bovine serum albumin (BSA) at 20 µg/mL as a control. After coating, wells were washed four times with 125 µL of 1% (w/v) BSA in PBS and then blocked for 1 hour at room temperature with 125 µL of 1% BSA in PBS. Immediately before the addition of bacteria, wells were washed six times with sterile PBS.

### Human host proteins-AbEnolase binding assay

Binding assays between human host proteins and purified *A. baumannii* enolase (AbEnolase) (Figure S1) were performed as previously described with some modifications^42^. *A. baumannii* Ab ATCC 17978 cells, grown overnight at 37°C in LB broth, were resuspended in PBS, centrifuged at 5000 *g* for 10 minutes, and washed twice with sterile PBS. Fifty microliters of increasing concentrations of AbEnolase (10, 50, and 100 mg/L) were added to wells pre-coated with plasminogen, fibronectin or fibrinogen, and incubated for 2 hours at room temperature. Wells were then rinsed six times with sterile PBS, and 50 µL of bacterial suspension was added to the wells for a 2-hours incubation to facilitate bacterial adhesion. Non-adherent bacteria were removed by washing the wells six times with sterile PBS. Adherent bacteria were collected by adding 125 µL of sterile PBS containing 0.5% Triton X-100. The lysates were diluted and plated onto LB agar, followed by incubation at 37°C for 24 hours to enumerate developed colonies. This allowed determination of bacterial adhesion to plasminogen, fibronectin and fibrinogen in the presence of AbEnolase.

### Human host proteins-bacteria-ENOblock binding assay

Human host proteins-bacteria-ENOblock binding assays were performed as described above with some modifications. The Ab ATCC 17978 strain was grown overnight at 37°C in LB, resuspended in PBS, and collected by centrifugation at 5,000× g for 10 min. The bacteria were washed twice in sterile PBS and resuspended in the same sterile buffer. A 50 µL of bacterial suspension was incubated with 0.5x and 1xMIC ENOblock for 30 min, then added to plasminogen-, fibronectin- or fibrinogen-coated wells and incubated 2 hours at room temperature for bacterial adsorption. The determination of bacterial adhesion to plasminogen, fibronectin and fibrinogen in the presence of ENOblock was performed as described above.

### *Galleria mellonella* infection model

*G. mellonella* infection model with Ab ATCC 17978 strain was established by haemocoel bacterial inoculation. Briefly, caterpillars obtained from Artroposfera (Toledo, Spain) were inoculated with 10 μL of the bacterial suspensions, which were incubated for 20-24 hours in LB at 37 °C. The minimal bacterial lethal dose 100 (MLD100) and LD50 were determined by inoculating various groups of larvae (8 *G. mellonella* per group) with decreasing amounts of Ab ATCC 17978 strains inocula from 10^6^ to 10^2^ CFU/mL, and monitoring the survival of the larvae for 7 days.

### Therapeutic efficacy of ENOblock in *G. mellonella* infection model

The efficacy of the ENOblock treatment was tested in *G. mellonella* survival assay as previously described^43^. Caterpillars were injected by the 10 μL of suspension containing MLD of *A. baumannii* Ab ATCC 17978. Treatment with 4xMIC of ENOblock was injected one hour post-infection. A group of larvae injected with 10 μL of sterile PBS was included as control. After inoculation, the larvae were incubated at 37°C in the dark and death was assessed over 3 days.

### Statistical Analysis

Group data are presented as means ± standard errors of the means (SEM). The student *t*-test was used to determine differences between means using the GraphPad Prism 9 (version 9.3.1; GraphPad Software, LLC.). For the *G. mellonella* survival model, a Kaplan-Meier test was performed to determine the difference between mortality rates. *P*<0.05 was considered significant.

## RESULTS

### High-throughput screening for repurposing drugs as antimicrobial agents

We developed and validated a high-throughput screen assay using the *A. baumannii* Ab ATCC 17978 and *E. coli* ATCC 25922 strains, as well as their respective MDR strains (**Figure 1A**). In total, we screened 2,464 compounds from the EU-OPENSCREEN ECBL Pilot library. We identified 33 compounds (1.32% of the total compounds) with inhibitory activity of ≥70% against at least one or both MDR strains of *A. baumannii* and *E. coli*. Of these 33 compounds, 7 showed MICs of ≤100 μM against the reference and MDR strains of *A. baumannii* and *E. coli* (**Figure 1A**). Among these 7 compounds, ENOblock (**Figure 1B**) was chosen for further studies. This compound was active against the *A. baumannii* Ab ATCC 17978 and *E. coli* ATCC 25922 strains, with AC50 values of 23.57 μM and 46.86 μM, respectively (**Figure 1C**).

**Figure 1:**
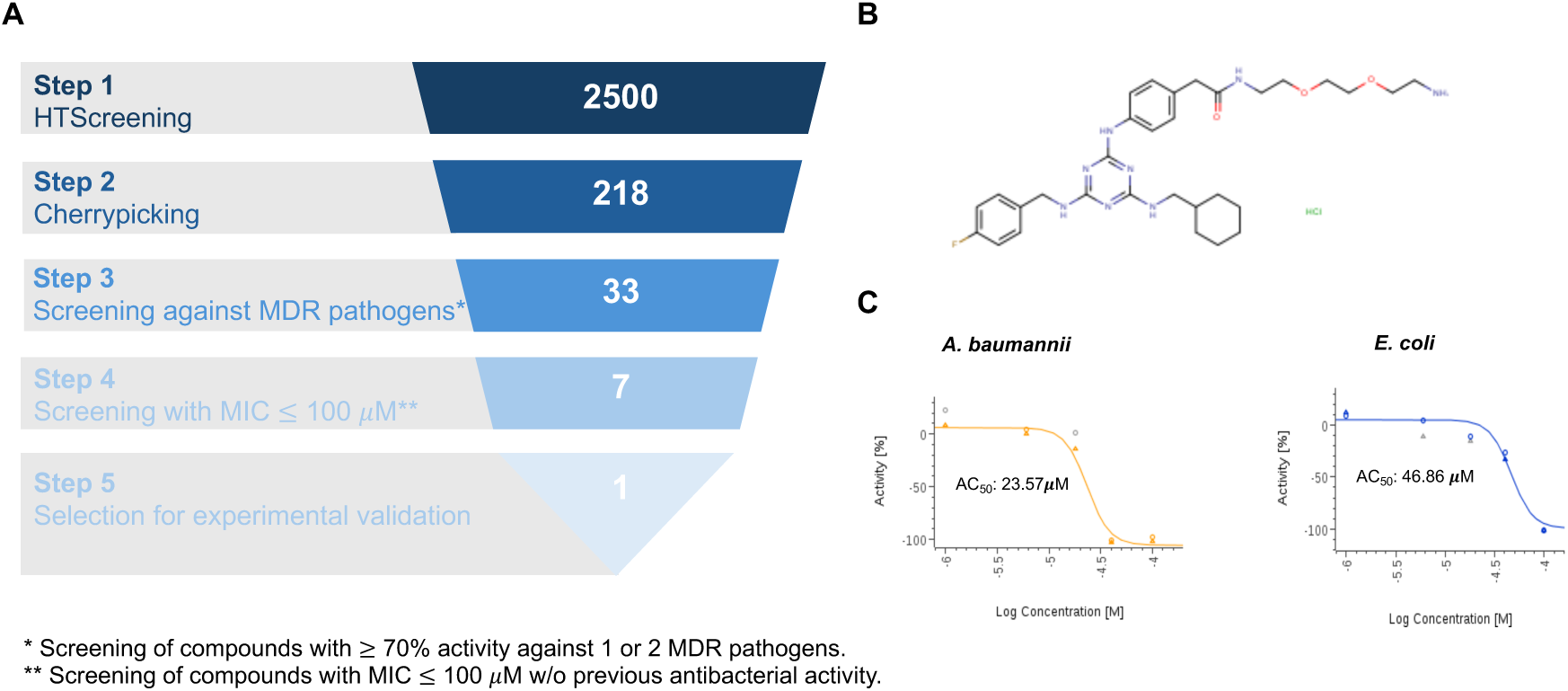
HTS in antibiotic discovery. (**A**) Hit identification workflow using the EU-OPENSCREEN library. *Screening of compounds with ≥ 70% activity against 1 or 2 MDR pathogens. **Screening of compounds with MIC≤100 µM w/o previous antibacterial activity. (**B**) Chemical structure of ENOblock. (**C**) Confirmation of ENOblock activity with the primary screening sample in dose-response mode against *A. baumannii* Ab ATCC 17978 and *E. coli* ATCC 25922 strains. AC50: Concentration at which 50% of maximum activity is observed.

### ENOblock is active against *A. baumannii*

To confirm the activity of ENOblock against clinical isolates of *A. baumannii*, ENOblock was tested against 14 and 18 colistin-resistant and carbapenem-resistant *A. baumannii* isolates. The results of the MICs tests are displayed in table 1 and table 2. The MICs ranged from 8 to 32 mg/L and 16 to 32 mg/L for ENOblock against colistin and carbapenem-resistant *A. baumannii*, respectively. The reference strain Ab ATCC 17978 present an ENOblock MIC of 8 mg/L. The MIC50 and MIC90 concentrations, which represent the concentration effective for 50 and 90% of the isolates tested, respectively, for ENOblock against colistin and carbapenems-resistant isolates were 16 and 32 mg/L, respectively. However, the MIC50 and MIC90 for colistin were 256 and >256 mg/L, and for carbapenems were 16 and 64 mg/l (**Tables 1 and 2**). Notably, the ENOblock MIC90 is three times lower than the CC50 of ENOblock in HeLa and macrophage cells (data not shown), and the selective pressure by growing Ab ATCC 17978 strain in the presence of increasing ENOblock concentrations did not allow *A. baumannii* to develop resistance to ENOblock (**Figure S2**). Using time-course assays, we evaluated the bactericidal activity of ENOblock against Ab ATCC 17978 and Ab CR17 strains. **Figure 2A** illustrates that ENOblock (2x and 4xMIC for Ab ATCC 17978 strain) exhibited bactericidal effect after 2, 4 and 8 hours, reducing the bacterial count by over 3 log10 CFU/mL compared to 0 hours. For the Ab CR17 strain, ENOblock (1x, 2x and 4xMIC for Ab CR17 strain) demonstrated a bactericidal effect after 2, 4 and 8 hours by reducing the bacterial count by over 3 log10 CFU/mL, compared to 0 hours. It is well known that the development of new repurposed drugs includes the assessment of the presence of synergy with clinically used antibiotics. To this end, we conducted a virtual screening of ENOblock in combination with different antibiotic families (colistin, imipenem, ceftazidime and tigecycline) using the EIIP/AQVN criterion to overcome bacterial resistance (**Table S2**). **Figure 2B** suggests that colistin and ENOblock with similar electronic properties, as indicated by their EIIP and AQVN values, tend to exhibit synergistic effects. Conversely, no synergistic activity was observed between the rest of antibiotics and ENOblock with different electronic properties. To confirm experimentally the *in silico* synergistic effect of ENOblock with colistin against *A. baumannii*, checkerboard and bacterial growth assays were performed (**Figure 2C, D**). Checkerboard assay indicated that ENOblock had a synergistic effect with colistin by enhancing the activity of colistin against Ab ATCC 17978 and Ab CR17 strains, resulting in an FIC index (FICI) of ≤0.5. In contrast, the combination of ENOblock with other antibiotics such as imipenem, ceftazidime and tigecycline did not increase their activities, yielding a FICI >0.5 (**Figure 2D**). Moreover, bacterial growth data confirms the checkerboard data showing that sub-MIC of colistin combined with ENOblock (8 mg/L) abolished completely the growth of Ab ATCC 17978 and Ab CR17 strains (**Figure 2E**).

**Table 1.** Antibacterial activity of ENOblock in colistin-resistant *A. baumannii* strains.

| Strains | Colistin MIC (mg/L) | ENOblock MIC (mg/L) |
| --- | --- | --- |
| #1 | 256 | 16 |
| #10 | >256 | 16 |
| #11 | 256 | 16 |
| #14 | 256 | 16 |
| #16 | >256 | 16 |
| #17 | 64 | 16 |
| #19 | >256 | 32 |
| #20 | >256 | 8 |
| #21 | 256 | 8 |
| #22 | >256 | 16 |
| #24 | 64 | 16 |
| #99 | >256 | 16 |
| #113 | >256 | 16 |
| Ab CR17 | 64 | 32 |
| MIC<sub>50</sub> | 256 | 16 |
| MIC<sub>90</sub> | >256 | 32 |

**Table 2.** Antibacterial activity of ENOblock in carbapenem-resistant *A. baumannii* strains.

| Strains | Imipenem/meropenem<br>MIC (mg/L) | ENOblock<br>MIC (mg/L) |
| --- | --- | --- |
| #17 | 8 | 16 |
| #37 | 16 | 16 |
| #40 | >64 | 16 |
| #53 | 64 | 16 |
| #286 | 16 | 16 |
| #288 | 32 | 16 |
| #289 | 8 | 16 |
| #295 | 8 | 16 |
| #298 | 16 | 32 |
| #299 | 16 | 16 |
| #405 | 32 | 32 |
| #410 | 32 | 32 |
| #414 | 32 | 32 |
| #416 | 32 | 32 |
| #417 | 16 | 16 |
| #440 | 8 | 16 |
| #441 | 8 | 16 |
| #448 | 8 | 32 |
| MIC<sub>50</sub> | 16 | 16 |
| MIC<sub>90</sub> | 64 | 32 |

**Figure 2.**
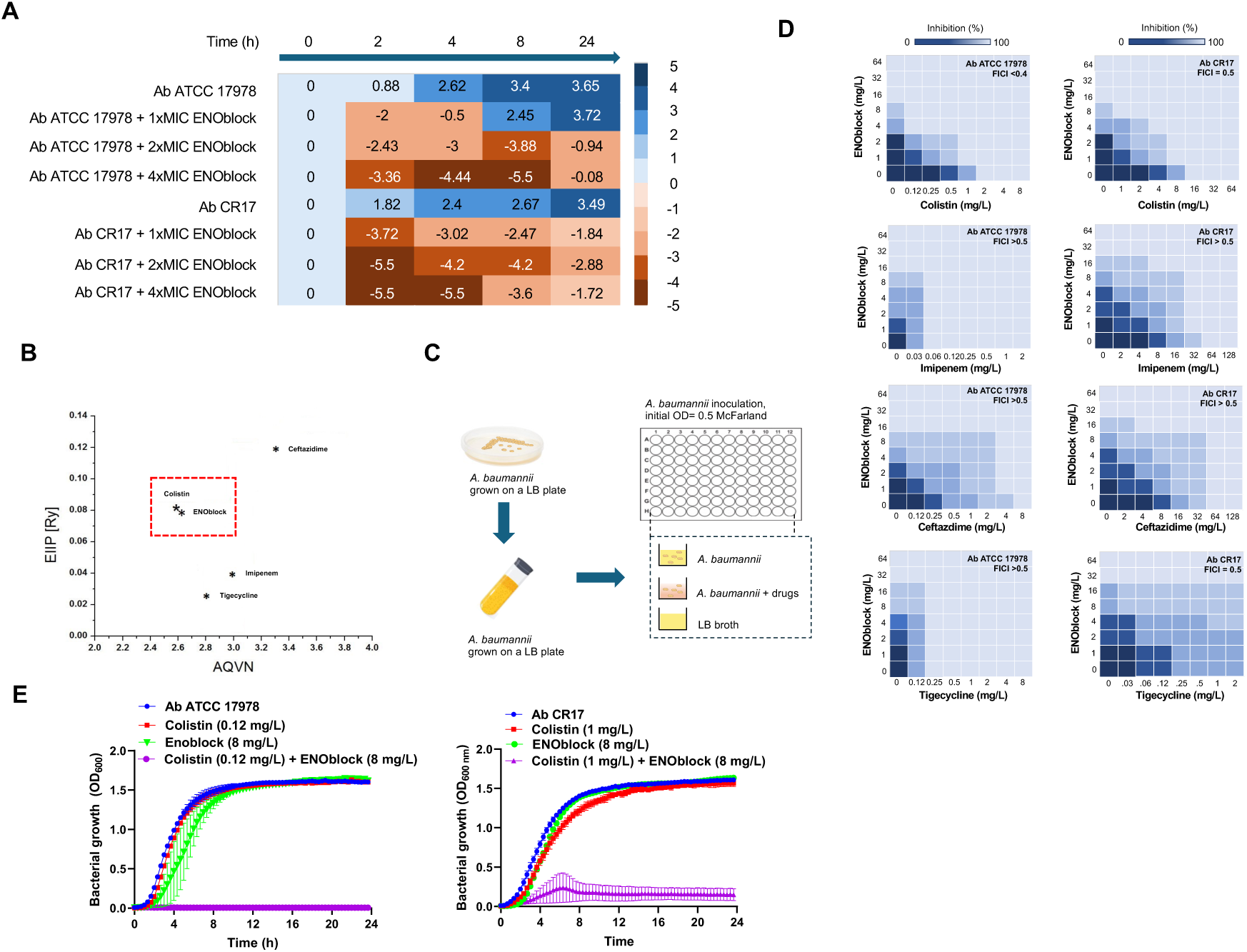
ENOblock is active against *A. baumannii* in monotherapy and in combination with colistin. (**A**) Time-kill curves of *A. baumannii* Ab ATCC 17978 and Ab CR17 strains in the presence of 1x and 1x, 2x and 4xMIC ENOblock for 24 hours. (**B**) Schematic presentation of the EIIP/AQVN criterion for the selection of ENOblock and colistin combination (green-penicillins, blue-carbapenems, red-quinolones). (**C** and **D**) Representative heat plots of microdilution checkerboard assay for the combination of ENOblock and colistin against Ab ATCC 17978 and Ab CR17 strains. (**E**) Bacterial growth curve plots of *A. baumannii* Ab ATCC 17978 and Ab CR17 strains in the absence and presence of colistin, ENOblock and colistin plus ENOblock. AQVN: Average quasi-valence number, EIIP: Electron-ion interaction potential.

### ENOblock inhibits the growth of *A. baumannii* via distinct mechanism of action

We employed the fluorescence microscopy-based BCP technique, as previously applied to *A. baumannii*^30,44,45^, to investigate the mechanism of action of ENOblock against Ab ATCC 17978 strain. The morphological changes induced by ENOblock were compared with those of antibiotics targeting major cellular pathways, including ciprofloxacin (DNA replication), rifampicin (RNA transcription), minocycline (protein translation), piperacillin and meropenem (cell wall synthesis), and colistin (membrane integrity). BCP results showed that ENOblock-treated cells exhibit unique morphological changes compared to those of antibiotic controls. In particular, membrane blebs were observed, as was the high SYTOX green signal of ENOblock-treated cells (**Figure 3A**), indicating the loss of membrane integrity. However, the ENOblock-treated cells showed different morphological changes from colistin-treated cells, a membrane integrity control, indicating that ENOblock interferes with bacterial membrane integrity but possibly in a manner different from colistin. Consistent with these differences, the image analysis profiles of ENOblock-treated cells clustered separately from those of untreated, other control antibiotics (**Figure 3B**). It is conceivable that ENOblock inhibits pathways that are distinct from those targeted by the comparator antibiotics, which collectively represent the most common modes of antibacterial action, in a manner similar to previous studies that have observed distinct profiles^35,46–48^. Altogether, the results showed that the ENOblock profile is distinct from the six antibiotic profiles.

**Figure 3.**
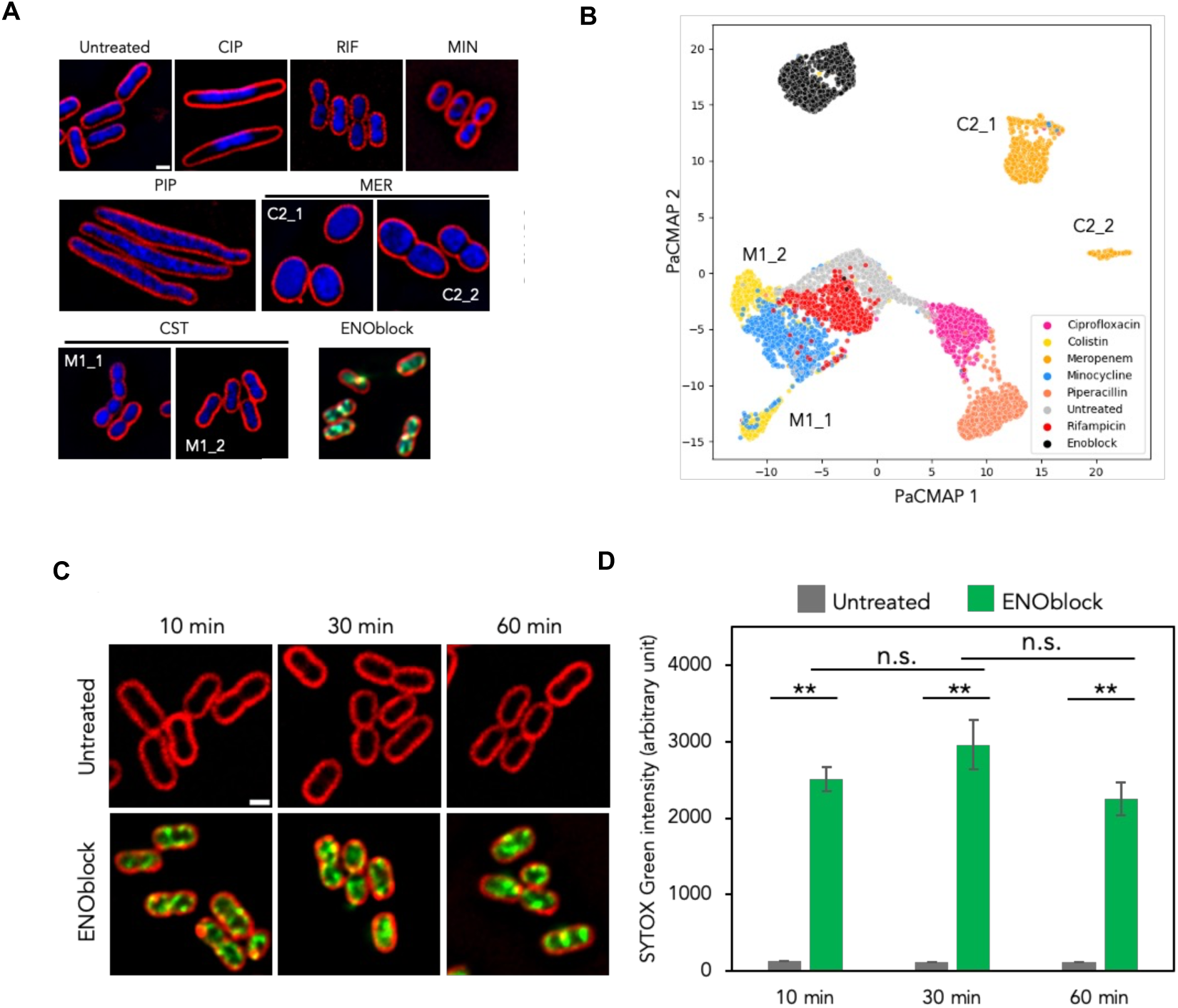
ENOblock exhibits antibacterial activity against *A. baumannii* via distinct mechanism of action and displays rapid permeabilization activity. (**A**) Representative images and (**B**) a PaCMAP plot of *A. baumannii* cells treated with 1x MIC of ENOblock and antibiotic controls. In all images, cell membranes are stained with FM4-64 (red), DAPI (blue), and SYTOX green (green). Scale bar represents 1 μM. (**C**) Representative images of *A. baumannii* cells treated with ENOblock for 10, 30 and 60 minutes, and then stained with FM4-64 (red) and SYTOX Green (green). Scale bar represents 1 μm. (**D**) A graph showing differences in SYTOX Green intensities between untreated and ENOblock-treated cells at different timepoints. Values represent mean intensities ± SEM of individual nucleoids from single cells, per condition. \*\**P*<0.01; two-tailed Student’s t-test, n.s.; not statistically significant.

### ENOblock exhibits rapid permeabilization activity against *A. baumannii*

The observation of a higher proportion of ENOblock-treated cells displaying high SYTOX Green intensity prompts us to consider the possibility of observing this impact at earlier time intervals, as membrane permeabilization often happens swiftly within minutes. Prior studies utilizing BCP demonstrated that a compound with the ability to disrupt membranes could impact the bacterial membrane in a mere 10 minutes timeframe^43,45^. Consequently, we conducted a temporal examination of SYTOX green staining on cells treated with ENOblock for 10, 30, and 60 minutes. The findings indicated that cells treated with ENOblock exhibited a notable rise in SYTOX intensity just 10 minutes after treatment (**Figure 3C, D**), in comparison to the control group that did not receive treatment (*P*<0.01). Although the fluorescence levels continued to increase after 30 minutes, this rise was not statistically significant (**Figure 3D**), indicating that the cells reached a saturation point with SYTOX Green after 10 minutes. Hence, the results indicates that ENOblock rapidly compromises the rigidity of the bacterial cell envelope, potentially resulting in cell death.

### ENOblock acts on *A. baumannii* through the inhibition of enolase

In order to shed light on the ENOblock mechanism of action, we docked ENOblock in the C-terminal domain of *A. baumannii* enolase. ENOblock exhibited a high docking score. The most stable pose shows that ENOblock binds to Ser371 and Asp207 amine acids through 3 hydrogen bonds (**Figure 4A**). Moreover, we quantified the interaction between ENOblock and purified AbEnolase (**Figure S1**) using an ITC assay. As shown in **Figure 4B**, ENOblock bound to AbEnolase with an affinity of 2.9 ± 0.78 µM. When Ser371 and Asp207 were together substituted with alanine, the binding of the ENOblock to the enolase is affected, making it impossible to calculate its KD constant, either because it decreases so much that the reaction do not reach the plateau on the binding curve (S371A) or because the protein loses stability causing its aggregation inside the cell (D207A and double mutant) as the ligand is added. (**Figure 4B and S3**). To investigate whether enolase in other Gram-negative bacteria, such as *E. coli* and *K. pneumoniae*, might also be a target of ENOblock, we performed *in silico* analyses to assess ENOblock’s binding capacity to enolase from both microorganisms and evaluated their susceptibility to ENOblock using a collection of MDR clinical isolates. Although the ENOblock binding sites in *E. coli* (GLN166, SER249, ASP245, ASP290, ASP316) and *K. pneumoniae* (ASP317, SER372) differed from the binding of some sites in *A. baumannii* (Asp207 and Ser371) (**Figure S4A, B**), ENOblock exhibited similar activity against MDR clinical isolates of *E. coli* and *K. pneumoniae*, with MIC50 values of 16 mg/L and 32 mg/L, respectively (**Tables S3 and S4**). As expected, ENOblock significantly reduced the growth of MDR *E. coli* Ec MCR1^+^ and *K. pneumoniae* Kp10 strains and increased their cell wall permeability (**Figure S5A, B**). To confirm that *A. baumannii* enolase is the potential target of ENOblock, we generated an enolase-deficient mutant. Deletion of the *enolase* gene in the Ab ATCC17978 strain (Δ*eno*) first abolished completely the enolase activity (**Figure 4C**), subsequently increased the ENOblock MIC from 8 to 32 mg/L (**Figure 4D**) and reduced the *A. baumannii* adherence to HeLa cells by >80% than the wt strain (**Figure 4E**). Furthermore, we examined the antibacterial activity of ENOblock against the Ab ATCC 17978 and Δ*eno* strains. **Figure 4F** reveals that the Ab ATCC 17978 strain exhibits rapid growth, reaching an OD of 1 within the first 4 hours. However, a significant disparity in growth is observed between the untreated cells and the cells treated with ENOblock, particularly at higher compound concentrations (16 and 32 mg/L). A different trend of growth inhibition is observed in the Δ*eno* strain, which shows a higher OD value compared with Ab ATCC 17978 strain in the presence of ENOblock treatment (**Figure 4G**). Similar results were observed with the *E. coli* enolase-deficient mutant (Ec Δ*eno*) strain (**Figure S6**). Considering these results, the difference in growth can be attributed to the resistance of the mutant strain to the ENOblock. The absence of enolase may hinder the compound’s ability to exert its effect, as indicated by the findings of the molecular docking study.

**Figure 4.**
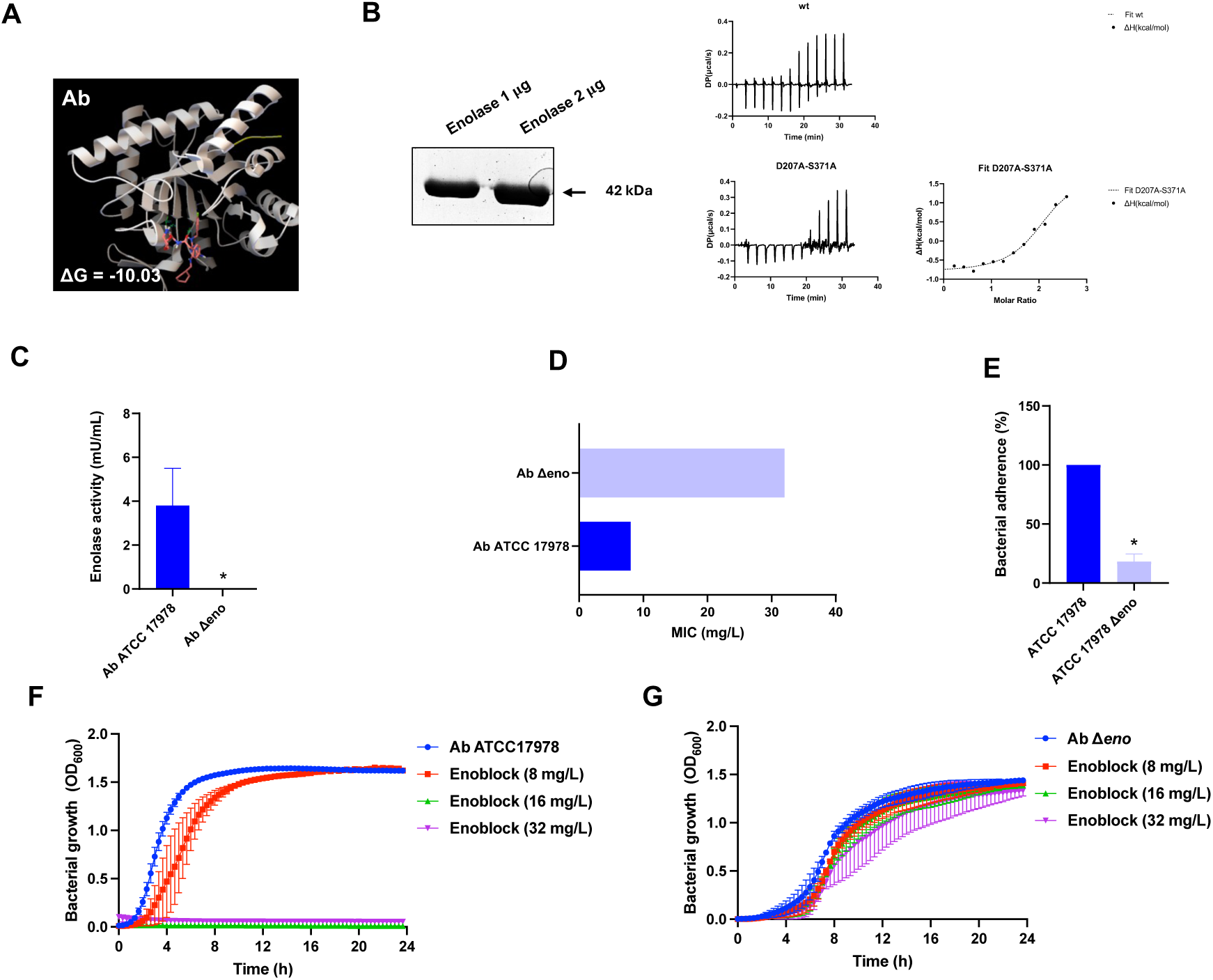
ENOblock acts on *A. baumannii* through the inhibition of enolase. (**A**) Structural models generated by docking of ENOblock into the C-domain of *A. baumannii* enolase. ENOblock is displayed as sticks. ΔG: Glide score. (**B**) Isothermal titration calorimetry (ITC) titrations with integrated fitted heat plots of ENOblock binding with wt enolase (wt) or enolase (D207A-S371A). (**C**) Enolase activity in determined in *A. baumannii* Ab ATCC 17978 and Δ*eno* strains. The data are presented as means ± SEM, and student t-test was used for statistical analysis. \**P* < 0.05, Ab ATCC 17978 vs Δ*eno*; two-tailed Student’s t-test. (**D**) MIC of ENOblock against *A. baumannii* Ab ATCC 17978 vs Δ*eno* strains. (**E**) *A. baumannii* Ab ATCC 17978 and Δ*eno* strains adhesion to HeLa cells. (**F** and **G**) Bacterial growth curve plots of *A. baumannii* Ab ATCC 17978 vs Δ*eno* strains in the absence and presence of ENOblock treatment at different concentrations. \**P* < 0.05, Ab ATCC 17978 vs Δ*eno*; two-tailed Student’s t-test.

### ENOblock affects the *A. baumannii*-host interaction

To evaluate the effect of ENOblock on the interaction between *A. baumannii* and host cells, we studied the adherence and invasion of the Ab ATCC 17978 and Ab CR17 strains on HeLa and macrophage cells for two hours in the presence of ENOblock (**Figure 5A**). We found that treatment with ENOblock at 1xMIC reduced the counts of adherent Ab ATCC 17978 and Ab CR17 strains on HeLa cells by 47% (*P*<0.05) and 31% (*P*<0.05), respectively, and on macrophage cells by 43% (*P*<0.05) and 29% (*P*<0.01) (**Figure 5B**). Notably, a more significant reduction was observed in the invasion of both strains, with ENOblock treatment reducing invasive counts in HeLa cells by 76% (*P*<0.01) and 46% (*P*<0.05), respectively, and in macrophage cells by 67% (*P*<0.01) and 41% (*P*<0.05), respectively (**Figure 5C**). Similarly, ENOblock at 1xMIC significantly reduced the adherence of *E. coli* MCR1^+^ and *K. pneumoniae* Kp10 to HeLa cells (**Figure S5C**). Furthermore, immunostaining of infected HeLa cells with the Ab ATCC 17978 and Ab CR17 strains, pretreated with ENOblock, showed a significant reduction in *A. baumannii* attachment to HeLa cells (**Figure 5D**).

**Figure 5.**
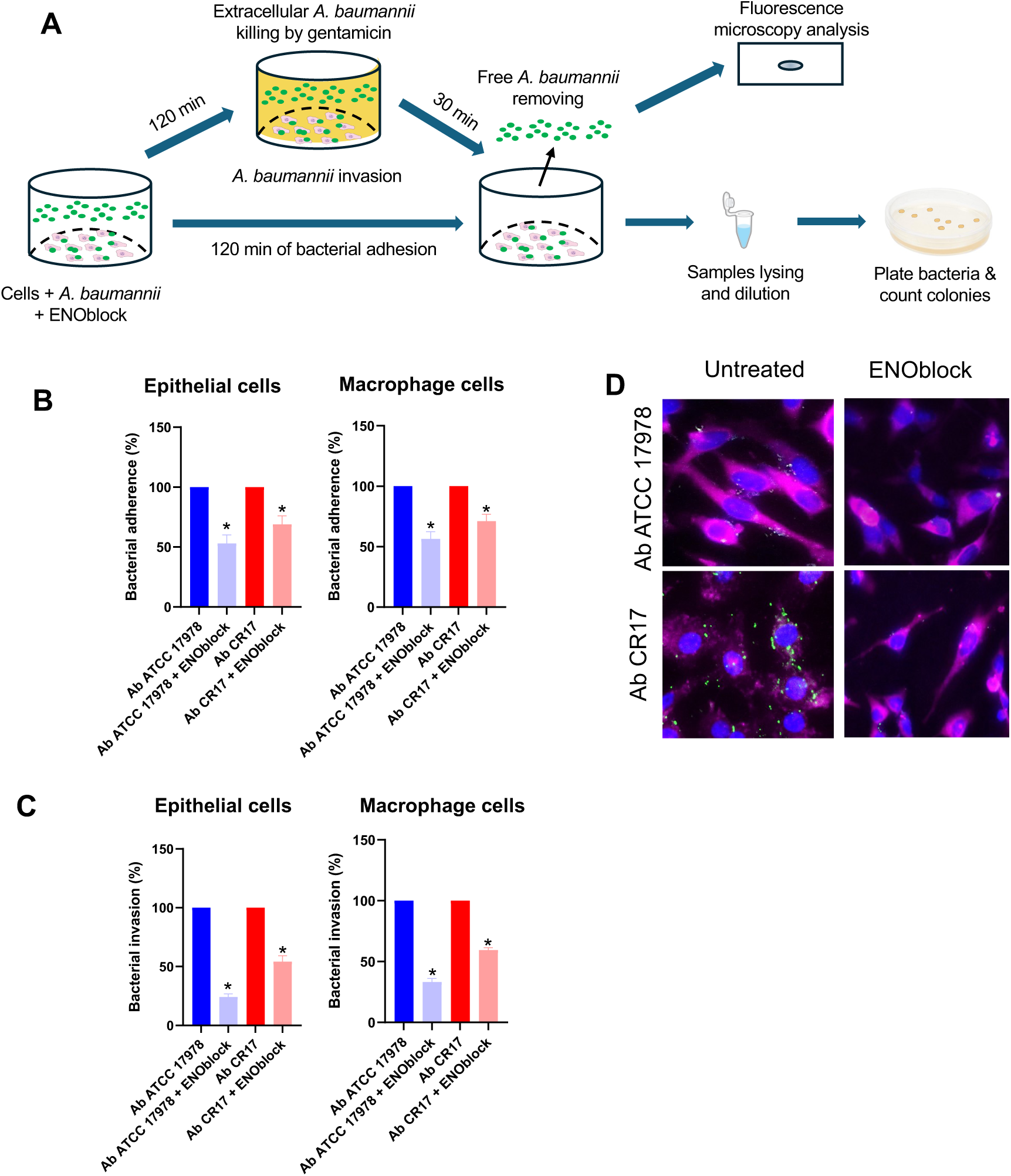
ENOblock affects the *A. baumannii*-host interaction. **(A)** Schematic of the bacterial adhesion/invasion assay. (**B** and **C**) Analysis of *A. baumannii* Ab ATCC 17978 and Ab CR17 strains adhesion/invasion into HeLa cells and macrophages with (1xMIC) and without ENOblock treatment. The data are presented as means ± SEM, \**P*<0.05: treatment vs no treatment; two-tailed Student’s t-test. (**D**) Immunostaining of fibronectin of HeLa cells (magenta) and Ab ATCC 17978 and Ab CR17 strains (green) pretreated with ENOblock (0x and 1xMIC), after bacterial adherence for two hours, was performed by specific primary antibodies against both strains and their respective secondary antibodies. Blue staining shows the location of HeLa cell nuclei.

### Host cell interaction induces metabolic changes in *A. baumannii* and upregulates enolase expression

Given that ENOblock reduces the interaction of *A. baumannii* with host cells and the Δ*eno* strain is defective in the adherence to host cells, we investigated whether enolase directly mediates this interaction. A TMT quantitative proteomics assay was performed on Ab ATCC 17978 strain alone and Ab ATCC 17978 strain in contact with epithelial and macrophage cells. Overall similarities and differences in protein expression between the two conditions (bacteria alone vs. bacteria in contact with host cells) were assessed using PCA (**Figure 6A**). As shown, Ab ATCC 17978 strain in contact with epithelial and macrophage cells displayed distinct protein expression patterns compared to bacteria alone, reflecting the impact of host cell interaction. Pairwise comparisons of differentially expressed proteins (DEPs) (bacteria in contact with epithelial cells *vs*. bacteria in contact with macrophage cells) revealed two key findings. First, overlapping patterns of overexpressed and downregulated proteins were observed (**Figure 6B**). Second, functional analysis of DEPs using Clusters of COG (**Figure 6C**) identified a common behaviour of Ab ATCC 17978 strain when is in contact with host cells. Overexpressed clusters were enriched for proteins involved in amino acid, inorganic ion, lipid, carbohydrate, and coenzyme transport and metabolism, transcription, defense mechanisms, post-translational modification, protein turnover, chaperones, and secondary metabolites. Notably, the number of significantly overexpressed proteins (red) exceeded that of downregulated proteins (blue) in response to bacterial interactions with epithelial and macrophage cells (**Figure 6D**). Enolase, the target of ENOblock, was significantly upregulated in *A. baumannii* upon contact with epithelial cells (log2 fold change > 2) and macrophage cells (log2 fold change > 1). These findings suggest that host cell interaction induces metabolic changes in *A. baumannii* and directly influences enolase expression.

**Figure 6.**
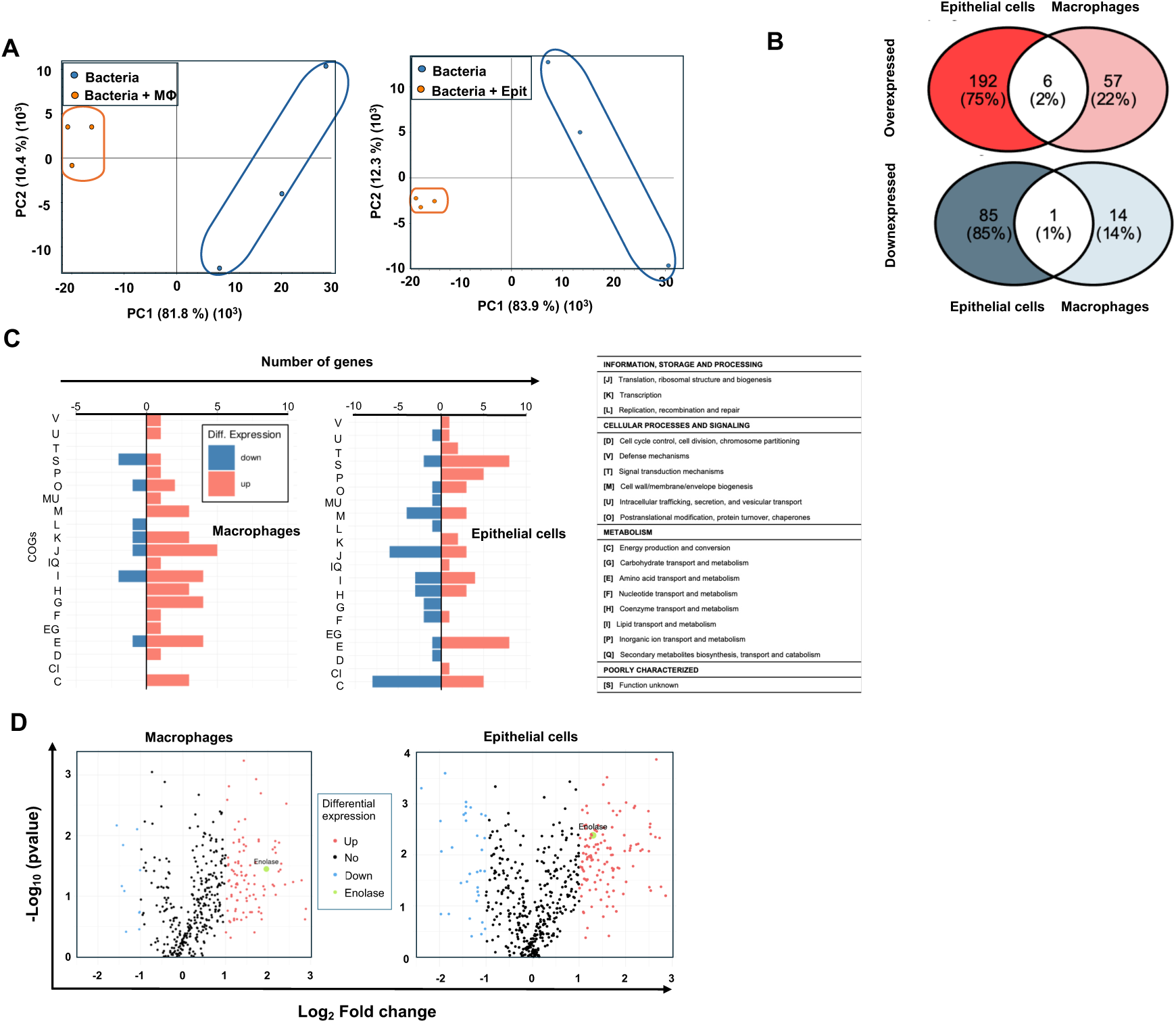
(**A**) PCA plot of TMT data from *A. baumannii* infected macrophages and epithelial cells (n = 3 for each group). **(B)** Venn diagrams showing the overlap of DEGs between the two comparisons: *A. baumannii* infected macrophages vs *A. baumannii* infected epithelial cells. The upper panel shows overlaps of overexpressed proteins, while the lower panel shows overlaps of downexpressed proteins. **(C)** COG category distribution of DEPs. The overexpressed and downexpressed proteins in each comparison are categorized by their COG functional groups, with blue representing downexpressed proteins and orange representing overexpressed proteins. **(D)** Volcano plots depicting the DEPs for the two comparisons. Proteins significantly overexpressed are shown in red, while significantly downexpressed proteins are shown in blue. Enolase overexpression is shown in green. Non-significant proteins are indicated in black. MΦ: macrophage cells, Epit: epithelial cells.

### Enolase mediates the interaction of *A. baumannii* with host cells via binding to host proteins

Given that *A. baumannii* upregulates enolase expression when in contact with host cells, and considering that enolase of *P. aeruginosa* and *Streptococcus suis* binds to human plasminogen and fibronectin on host cells, respectively^49,50^, we sought to determine whether *A. baumannii* enolase binds to host proteins (Plasminogen, fibronectin and fibrinogen) to mediate host cell interactions with *A. baumannii*. ISM analysis and purified AbEnolase (**Figure S1**) were used to assess interactions with host proteins. As shown in **Figure 7A-C**, ISM-SM was employed to identify shared informational characteristics among *A. baumannii* enolase, host proteins (Plasminogen, fibronectin and fibrinogen), and ENOblock. The ISM is based on the principle that interacting proteins and small molecules share common informational properties, appearing as cross-spectral peaks^42^. Our consensus spectral analysis revealed a characteristic peak at frequency F(0.270) shared by plasminogen, enolase and ENOblock (**Figure 7A**). The same peak was observed when fibronectin, enolase and ENOblock were analyzed (**Figure 7B**). However, a not primary frequency peak F(0.435) has been shared by fibrinogen, enolase and ENOblock (**Figure 7C**). These findings suggest that shared informational content at F(0.270) and F(0.435) indicates potential interactions between enolase and human proteins. In the next step, the plasminogen, fibronectin or fibrinogen – enolase complexes were modelled. Obtained sequences of enolase and plasminogen, fibronectin or fibrinogen were sent to Alfafold3 to build a protein-protein complex. The intermolecular interactions from the obtained complex were according to ISM method–detected interaction domains (**Figure 7E-G**, yellow ribbons). Finally, enoblock was docked into the plasminogen-enolase cavity, fibronectin-enolase cavity and fibrinogen-enolase cavity, using Autodock Vina software. The obtained confirmation had a docking energy of -9.3, -7.9 and -9.9 kcal/mol, respectively. To experimentally validate these observations, we first performed binding assays using purified AbEnolase with ECM proteins. The adhesion of Ab ATCC 17978 strain to immobilized plasminogen, fibronectin or fibrinogen was significantly reduced with the addition of increasing concentrations of AbEnolase or ENOblock (**Figure 7G-L**). However, the incubation of Ab ATCC 17978 strain with 0.5x and 1xMIC during 2 hours did not reduce the bacterial concentration (data not shown).

**Figure 7.**
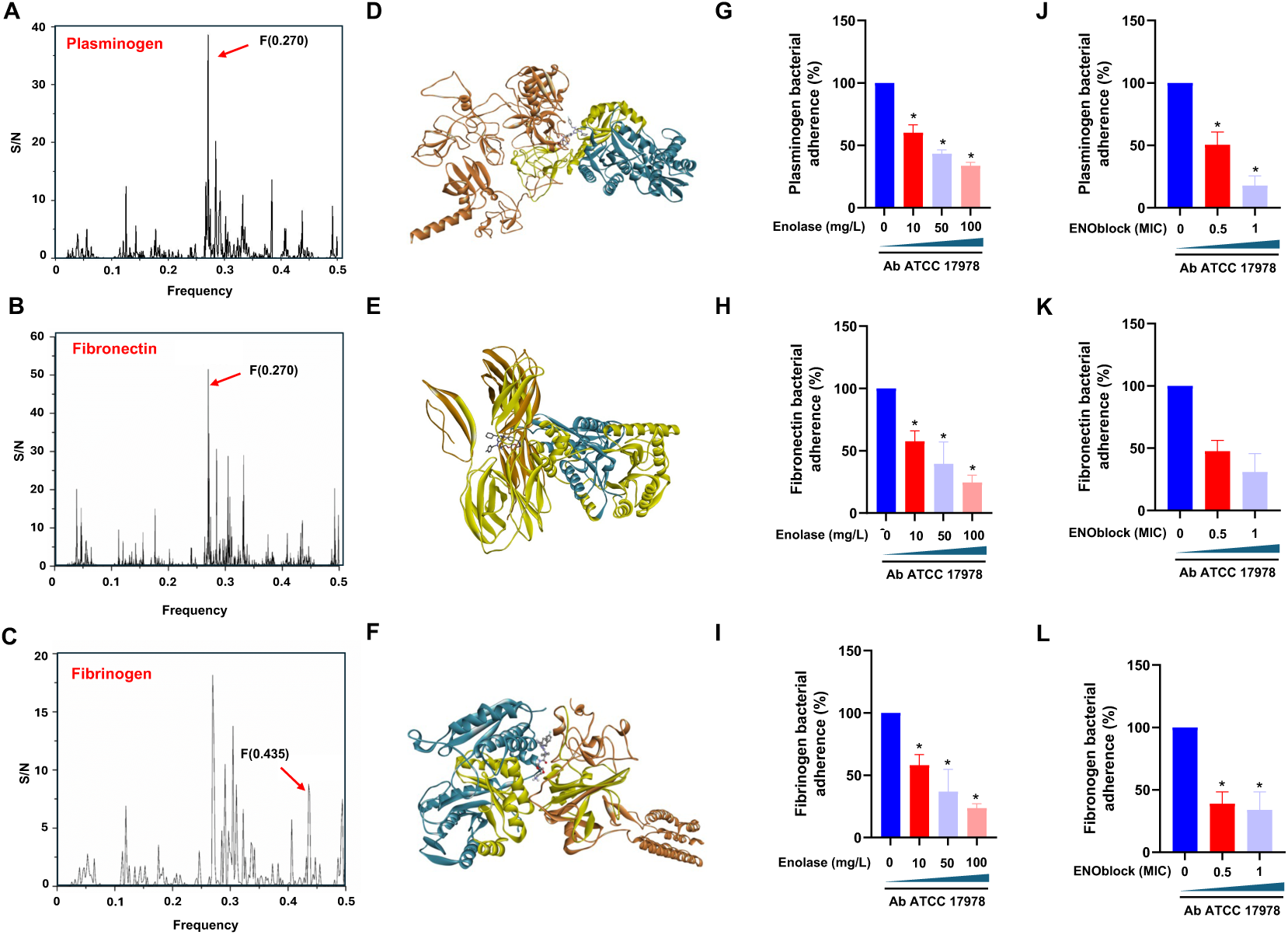
Enolase mediates the interaction *A. baumannii* of host cells via binding to host proteins. (**A-C**) Consensus-spectrum (CIS) of enolase, plasminogen and ENOblock, enolase, fibronectin and ENOblock, and enolase, fibrinogen and ENOblock (**D-F**) Structural models of protein-protein complex generated by Alfafold3 analysis by docking of ENOblock (ball and stick), enolase (pelorous ribbons) and plasminogen, fibronectin or fibrinogen (tangerine ribbons). The intermolecular interactions from the obtained complex were according to ISM method–detected interaction domains (yellow ribbons). (**G-I**) Inhibition of *A. baumannii* adherence to immobilized plasminogen, fibronectin and fibrinogen by free enolase. Ab ATCC 17978 strain was incubated in plasminogen, fibronectin and fibrinogen-coated wells for 3 h at room temperature, containing increasing concentrations of free enolase (0, 10, 50 and 100 mg/L). (**J-K**) Inhibition of *A. baumannii* adherence to immobilized plasminogen, fibronectin and fibrinogen by ENOblock. Ab ATCC 17978 strain was incubated in plasminogen, fibronectin and fibrinogen-coated wells for 2 h at room temperature, containing increasing concentrations of ENOblock (0.5x and 1xMIC). Adherent bacteria to plasminogen, fibronectin and fibrinogen-coated wells were quantified by serial dilutions as described in <u>materials and methods</u>. Results were expressed as the percentage of total untreated *A. baumannii* adhered to immobilized plasminogen, fibronectin and fibrinogen.

### ENOblock presents therapeutic efficacy *in vivo*

To confirm the *in vitro* effect of ENOblock in monotherapy and in combination with colistin against *A. baumannii*, and to study this efficacy in a complete organism, we moved to an invertebrate model of infection by *A. baumannii*. First, we determined the virulence of the Ab ATCC 17978 strain after haemocoel administration in *G. mellonella* (**Figure 8A**). The mortality rates of animals were inoculum concentration dependent. LD50 and MLD100 for the Ab ATCC 17978 strain were 10^2^ and 1x10^5^ CFU/mL, respectively (**Figure BC**). Subsequently, in a *G. mellonella* model of infection, we administered ENOblock (32 mg/L) to animals after haemoceol administration of an MLD100 of the Ab ATCC 17978 strain (**Figure 8C**). Animals receiving treatment with ENOblock showed significantly a greater increase in survival compared with untreated animals (*P*<0.01) (**Figure 8D**).

**Figure 8.**
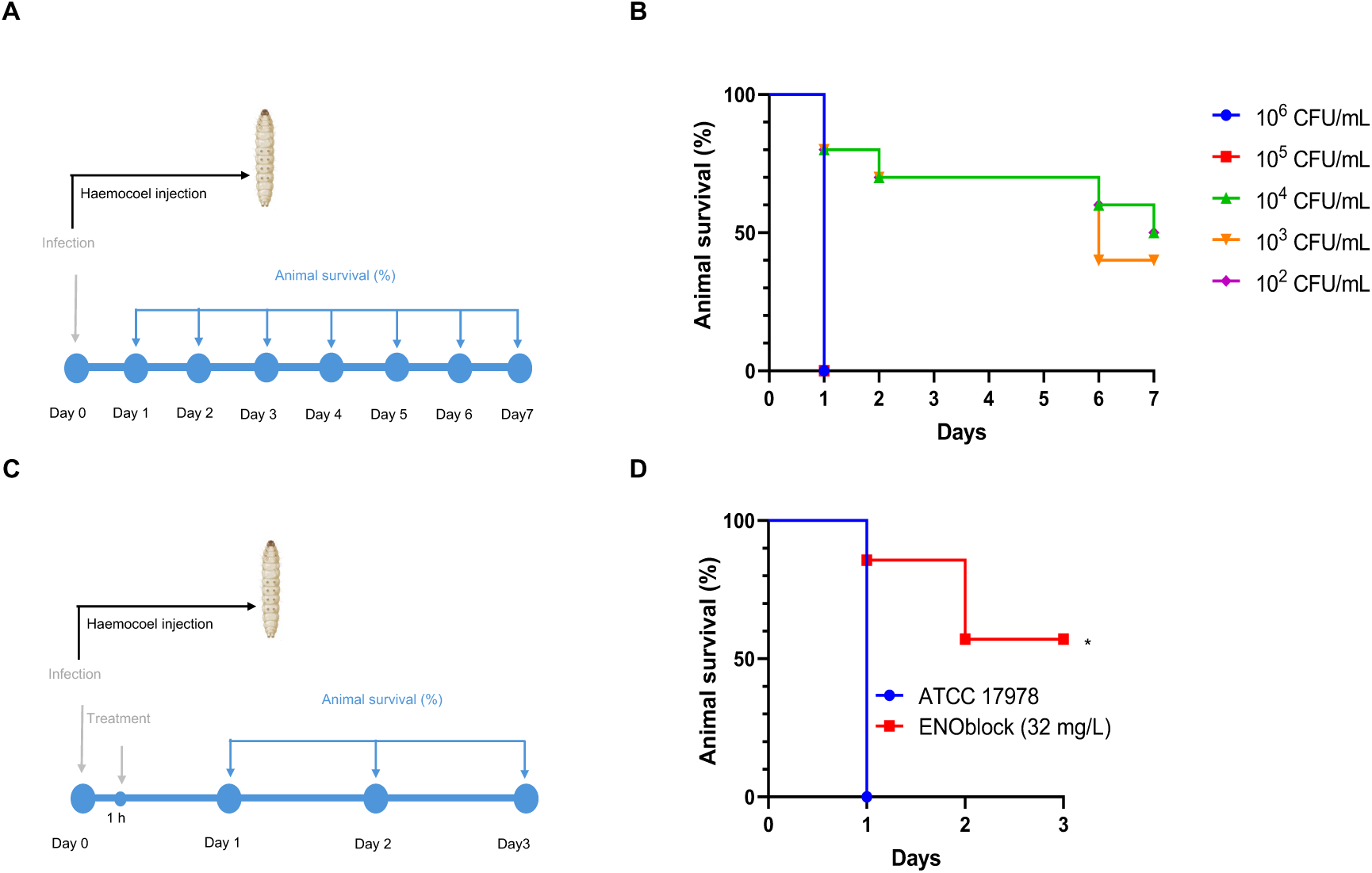
ENOblock displays efficacy in *G. mellonella* model of infection. (**A**) Experimental design for *A. baumannii* Ab ATCC 17978 strain lethal doses determination. (**B**) Seven days of mortality monitoring in *G. mellonella* administered with different inoculum of *A. baumannii* Ab ATCC 17978 strain (n = 8 per group). (**C**) Experimental design for *A. baumannii* Ab ATCC 17978 strain infection and treatment. (**D**) Three days of mortality monitoring in *G. mellonella* administered with MDL100 (10^5^ CFU/mL) *A. baumannii* Ab ATCC 17978 and treated or not with ENOblock (32 mg/L; n = 8 per group). *P*<0.01, treatment vs no treatment; Kaplan-Meier test.

## DISCUSSION

Due to the antibacterial effects of various anticancer drugs on *A. baumannii*^51–54^, we hypothesized that ENOblock, identified in this study after a HTS of the EU-OPENSCREEN library, might exhibit strong antibacterial activity against colistin- or carbapenem-resistant *A. baumannii*. After initial antimicrobial confirmation of ENOblock, we additionally tested its susceptibility against 32 clinical isolates of *A. baumannii* resistant to colistin or carbapenems (**Tables 1 and 2**). The ENOblock MIC90 is 32 mg/L, which is two to more than six times lower than the MIC90 of carbapenem and colistin, respectively. This MIC90 value falls within the range of other known antibiotics such as amikacin, amoxicillin-clavulanic acid, ceftazidime-avibactam, and fosfomycin, among others^55^. Additionally, it is three times lower than the CC50 observed in epithelial and macrophages cells. This suggests that ENOblock, at MIC90 concentration, is unlikely to be toxic or cause off-target effects in humans, despite targeting enolase, a highly conserved enzyme also present in human cells.

It is noteworthy that ENOblock demonstrated similar activity against MDR *E. coli* and *K. pneumoniae*, consistent with previously published data on the antibacterial activity of the anticancer drug family tamoxifen and its metabolites used in earlier studies^52–54^. Specific AQVN/EIIP domains, combined with structural properties, can serve as effective filters for the virtual screening of molecular libraries to identify new drug candidates, including new antibiotics. Using molecular descriptors, the EIIP and AQVN we have proposed suitable antibiotics for treating MDR bacterial infections^56^. In this study, we analyzed the electronic properties of ENOblock and antibiotics to which Ab ATCC 17978 is sensitive. Our analysis suggests that antimicrobials with similar electronic properties tend to act synergistically. However, the limited number of molecules analyzed restricts our ability to establish a criterion for predicting synergy in *A. baumannii*. Moreover, studies show that the molecular mechanisms underlying such synergistic effects remain not fully understood^57^. Nonetheless, the presented results with the observed tendency for ENOblock and colistin, to exhibit synergy suggest that similar electronic properties may contribute to effective antibacterial combinations. Small molecules with similar AQVN and EIIP values have previously been shown to interact with the common therapeutic target^40,58^.

The antibacterial activity of ENOblock at 1x, 2x, and 4xMIC against the colistin-resistant strain (Ab CR17) is higher than against the colistin-susceptible reference strain (Ab ATCC 17978). This result may be related to differences in the cell wall structure of the two strains, where colistin-resistant strains of *A. baumannii* are more permeable than colistin-susceptible strains^24,59,60^.

The antimicrobial activity of ENOblock identified in this study suggests promising potential that warrants further exploration *in vivo* after determining its pharmacokinetic parameters. However, *in vitro* bacterial growth showed a progressive regrowth of the Ab ATCC 17978 strain after treatment with ENOblock at 1xMIC, suggesting that this strain may have acquired resistance to this compound. It is worth noting that the MIC of ENOblock against the Ab ATCC 17978 strain in these time-kill assay conditions is 8 mg/L, which is below the 2x and 4xMIC of ENOblock. Further investigations, including the determination of its concentration during the time-kill assay, are necessary to better understand the regrowth of this strain in the presence of ENOblock.

As with other antimicrobial agents, there is a risk of emerging resistance. *A. baumannii* adaptation mechanisms could lead to reduced susceptibility to ENOblock, either through mutations in enolase or compensatory metabolic pathways. However, selective pressure against *A. baumannii* did not induce resistance to ENOblock. Consistent with this data, combination therapies, which are known to mitigate the risk of resistance development, have shown synergy between ENOblock and colistin.

Additionally, BCP analysis showed that ENOblock-treated *A. baumannii* exhibits distinct morphological changes compared to comparator antibiotics (**Figure 3A, B**), suggesting that ENOblock inhibits pathways different from those targeted by the comparator antibiotics. Membrane-disruptive agents can be classified into various subcategories, each exhibiting a unique structure on BCP. Nevertheless, the BCP profile of ENOblock-treated cells observed in this study did not precisely correspond to any previously described profiles^35,44–47^. Consequently, the specific target on the membrane for ENOblock could not be determined, but the presence of SYTOX Green indicates that ENOblock can permeabilize membranes. Therefore, future studies should investigate the precise locations on the bacterial membrane where ENOblock is active.

It is widely known that ENOblock inhibits enolase activity in eukaryotic cells^61^, and enolase, a cytoplasmic glycolytic enzyme, is a key component of the RNA degradosome in Gram-negative bacteria such as *E. coli* and *P. aeruginosa*^62^. For this reason, we decided to conduct a preliminary computational study to determine if ENOblock could potentially bind to enolase in *A. baumannii*. As a result, ENOblock exhibited a better docking score. Deletion of *enolase* gene in *A. baumannii* increased the ENOblock MIC four-fold and increased bacterial growth in the presence of ENOblock, suggesting that enolase could be a potential target of ENOblock in *A. baumannii*. The interaction between AbEnolase and ENOblock was confirmed with a microcalorimetric assay where recombinant expressed and purified AbEnolase showed a *KD* 2.9 µM (**Figure 4B**). The computational study showed two potential aminoacids (D207 and S371) in close proximity in the predicted 3D structure that could be part the binding site of ENOblock. To confirm this hypothesis, AbEnolase with both single and the double mutations were recombinantly expressed and purified, where the aminoacids of interest were exchange for an alanine. These mutants showed a decrease in affinity against the ENOblock. The mutation in the Serine 370 caused a reduction in the affinity while the mutation in the Aspartic acid 207 led not only to decreased affinity but also to decreased protein stability causing protein aggregation from a ligand:enolase Molar Ratio of 1.5. The double mutation showed a stronger effect in protein stability and ligand affinity decrease.

Importantly, two studies have reported that enolase also resides on the cell wall outer membrane of *E. coli* and *P. aeruginosa*^49,63^ and mediates the binding of *P. aeruginosa* and *S. suis* to plasminogen and fibronectin on host cells^49,50^. In this study, we showed that ENOblock reduces the interaction of *A. baumannii* with host cells, possibly by affecting the host proteins binding activity of *A. baumannii* enolase. Other components of the *A. baumannii* cell wall such as Tuf, OmpA and chaperone-usher pathway pili binds also to plasminogen, fibronectin and fibrinogen, respectively^42,64,65^. Consequently, the possibility of multi-targeting, with ENOblock binding to both cytosolic and membranal enolase, might contribute to its overall activity, which is an attractive aspect of ENOblock’s antibacterial properties.

We have demonstrated through various assays that ENOblock can be repurposed as an antibacterial agent. However, its *in vitro* efficacy might not fully translate to *in vivo* conditions due to differences in pharmacokinetics, tissue distribution, and metabolic stability. Our preliminar *in vivo* experiments have shown that ENOblock, at 32 mg/L, has been able to increase animal survival in presence of *A. baumannii*. This result aligns with previous observations that deletion of *enolase* gene abolishes the virulence of *P. aeruginosa* in a murine acute pneumonia model^66^. Further *in vivo* studies and preclinical models are necessary to confirm the therapeutic potential and safety profile of ENOblock, and future work should focus on detailed toxicity assessments and the evaluation of ENOblock’s selectivity for bacterial versus human enolase.

In summary, this drug discovery approach ought to be viewed as the first step in creating a brand-new class of antimicrobial drugs. The efficacy of ENOblock can be investigated further by combining this newly repurposed compound with clinically used antibiotics (like colistin) in *in vivo* experiments, with the aim of reducing mortality and improving therapeutic efficacy in cases of severe infections, even with the current antimicrobial therapies.

## Acknowledgments

This research was funded by the Ministerio de Ciencia e Innovación, Agencia Estatal de Investigación, Fondo Europeo de Desarrollo Regional, MCIN/AEI/10.13039/501100011033/FEDER, UE (Grant PID2022-136357OBI00), and (Grant CEX2020-001088-M-20-5), by the Consejería de Universidad, Investigación e Innovación de la Junta de Andalucía (Grant ProyExcel_00116), by the National Research Council of Thailand (NRCT) and Mahidol University: N42A650368, and by the Ministry of Science, Technological Development and Innovation of the Republic of Serbia (Grant number 451-03-136/2025-03/200017). We acknowledge EU-OPENSCREEN ERIC for providing its compound collection and Fundación MEDINA HTS antimicrobial screening platform to support the discovery of the antibacterial activity of the compound described in the presented work. This article is based upon work from COST Action EURESTOP, CA21145, supported by COST (European Cooperation in Science and Technology).

A.M.R is supported by a doctoral fellowship PRE2022-104318, from the Agencia Estatal de Investigación, Ministerio de Ciencia e Innovación.

## Conflict of interest

The authors declare that the research was conducted in the absence of any commercial or financial relationships that could be construed as a potential conflict of interest.

## Author contributions

I.M.P., A.M.R., M.C., L.T.G, P.S., T.S., M.S. & A.H. methodology, investigation, formal analysis. A.P.P, S.G., O.G. A.H. & P.N. writing-review and editing, methodology, investigation. Y.S. writing-review and editing, supervision, funding acquisition, conceptualization.

## Data availability statement

The data supporting the findings of this study are available from the corresponding author upon reasonable request.

## Supplementary data

**Table S1.**
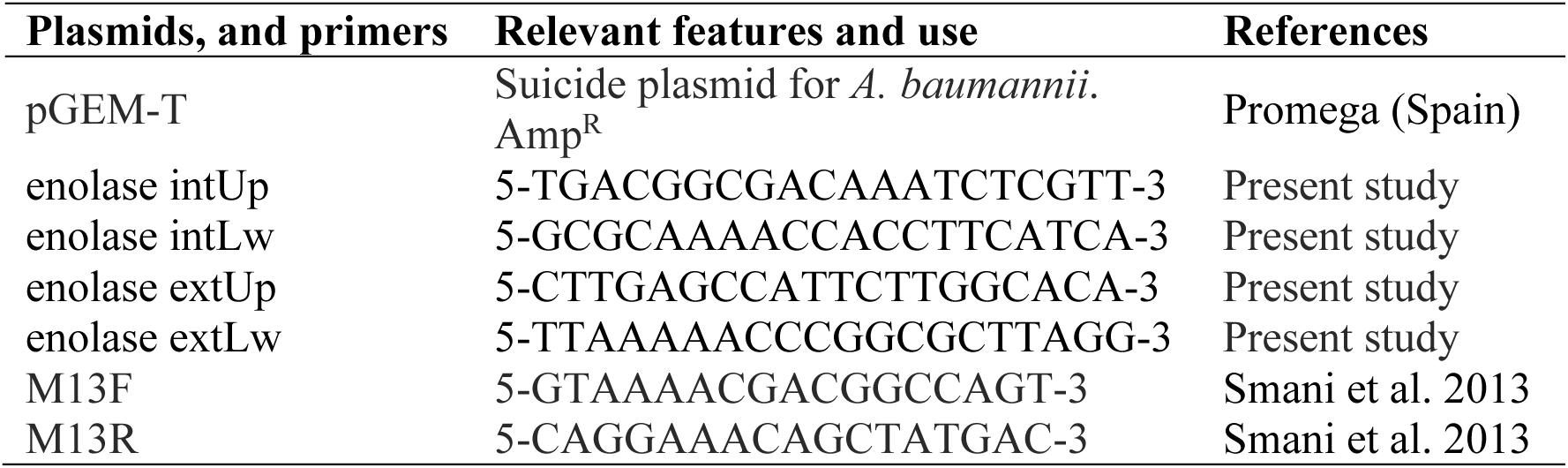
Plasmids and primers used in this study.

**Table S2.** EIIP and AQVN parameters to analyze the electronic properties of drugs tested for synergy against *A. baumannii* Ab ATCC 17978 strain.

| **Molecule** | **AQVN** | **EIIP** |
| --- | --- | --- |
| ENOblock | 2.628 | 0.078 |
| Colistin | 2.592 | 0.084 |
| Imipenem | 2.973 | 0.034 |
| Ceftazidime | 3.288 | 0.127 |
| Tigecycline | 2.815 | 0.025 |

AQVN: and average quasi-valence number, EIIP: Electron-ion interaction potential.

**Table S3.** Antibacterial activity of ENOblock in colistin-resistant *E. coli* strains.

| **Strain#** | **Colistin resistance mechanism** | **Colistin MIC (mg/L)** | **ENOblock MIC (mg/L)** |
| --- | --- | --- | --- |
| **CRA5** | - | 8 | 16 |
| **CRA7** | - | 8 | 16 |
| **CRA8** | MCR1 | 8 | 16 |
| **CRA17** | - | 8 | 32 |
| **CRA20** | MCR1 | 8 | 32 |
| **CRA32** | - | 8 | 8 |
| **CRA57** | - | 4 | 16 |
| **R2** | MCR1 | 8 | 32 |
| **R5** | MCR1 | 4 | 32 |
| **R6** | MCR1 | 8 | 16 |
| **R34** | MCR1.5 | 8 | 64 |
| **R37** | MCR3.2 | 8 | 32 |
| **MCR1+** | MCR1 | 4 | 64 |

**Table S4.** Table 1. Antibacterial activity of ENOblock in carbapenem-resistant *K. pneumonaie* strains.

| **Strain#** | **Meropenem**<br>**resistance**<br>**mechanism** | **Meropenem**<br>**MIC (mg/L)** | **ENOblock**<br>**MIC (mg/L)** |
| --- | --- | --- | --- |
| **Kp6** | KPC-3 | 2 | 32 |
| **Kp7** | KPC-2 | - | 16 |
| **Kp8** | KPC-2 | - | 64 |
| **Kp9** | KPC-41 | - | 64 |
| **Kp10** | KPC-2 | 16 | 32 |
| **Kp11** | KPC-2 | 4 | 16 |
| **Kp12** | KPC-2 | 16 | 64 |
| **Kp13** | KPC-3 | >16 | 64 |
| **Kp14** | KPC-2 | 4 | 64 |
| **Kp15** | KPC-11 | 4 | 64 |
| **Kp16** | KPC-3 | >16 | 64 |
| **Kp17** | KPC-3 | 16 | 16 |
| **Kp18** | KPC-2 | 4 | 32 |
| **Kp19** | KPC-3 | 4 | 64 |

**Figure S1.**
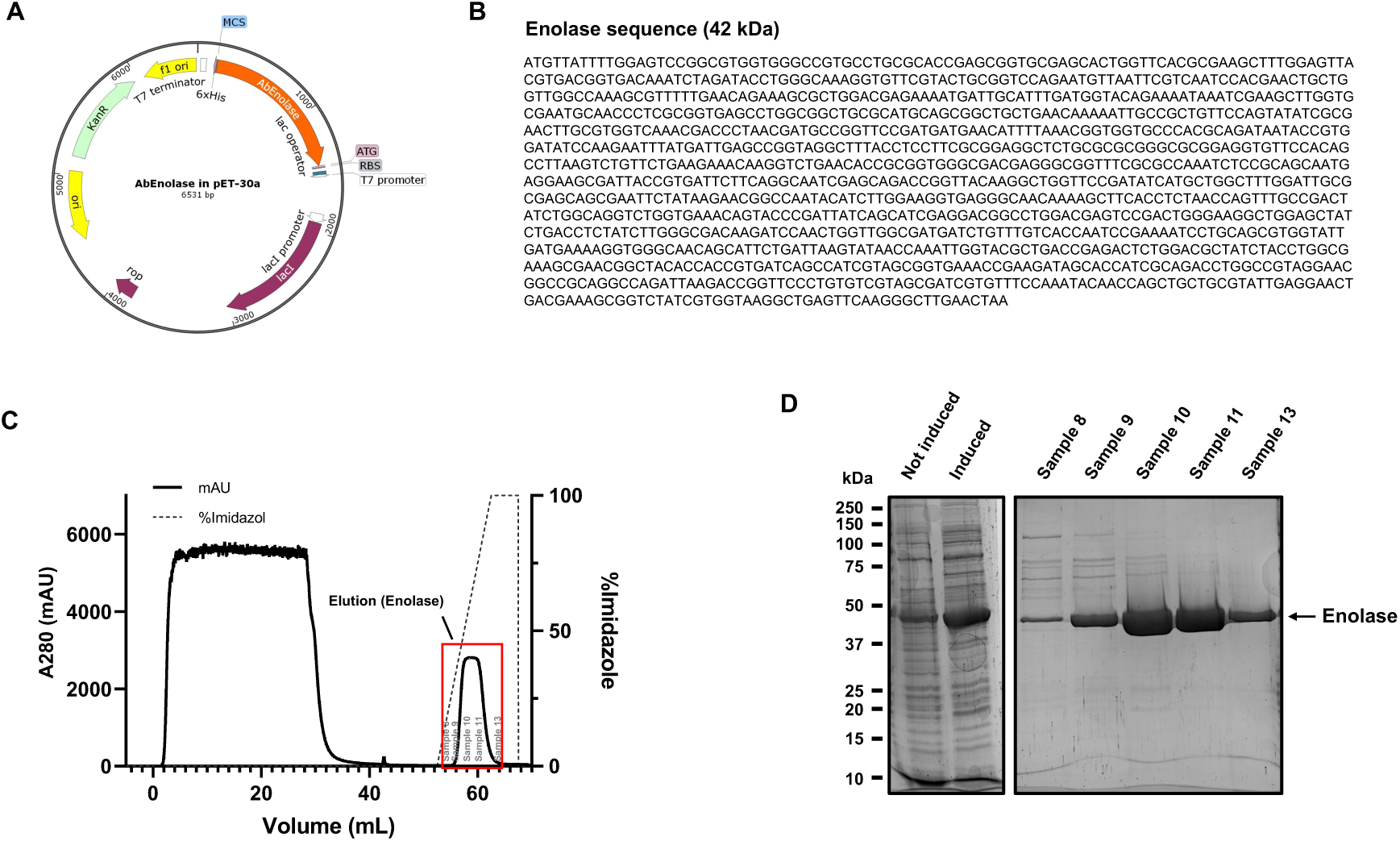
Construction and characterization of *A.baumannii* enolase. **(A)** Illustration of the recombinant expression vector of enolase. **(B)** The sequence of the enolase. **(C)** Purification of enolase using the Histrap FF column. **(D)** SDS-PAGE gel analysis of the purified enolase.

**Figure S2.**
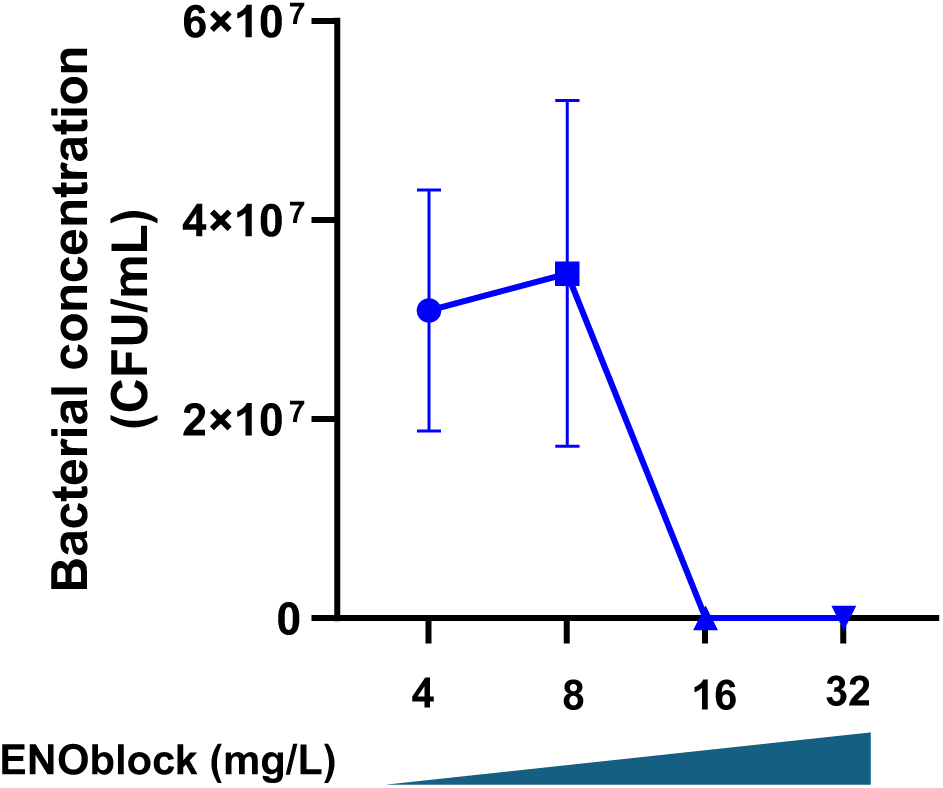
ENOblock selective pressure. *A. baumannii* Ab ATCC 17978 concentrations after incubation with increasing concentrations of ENOblock.

**Figure S3.**
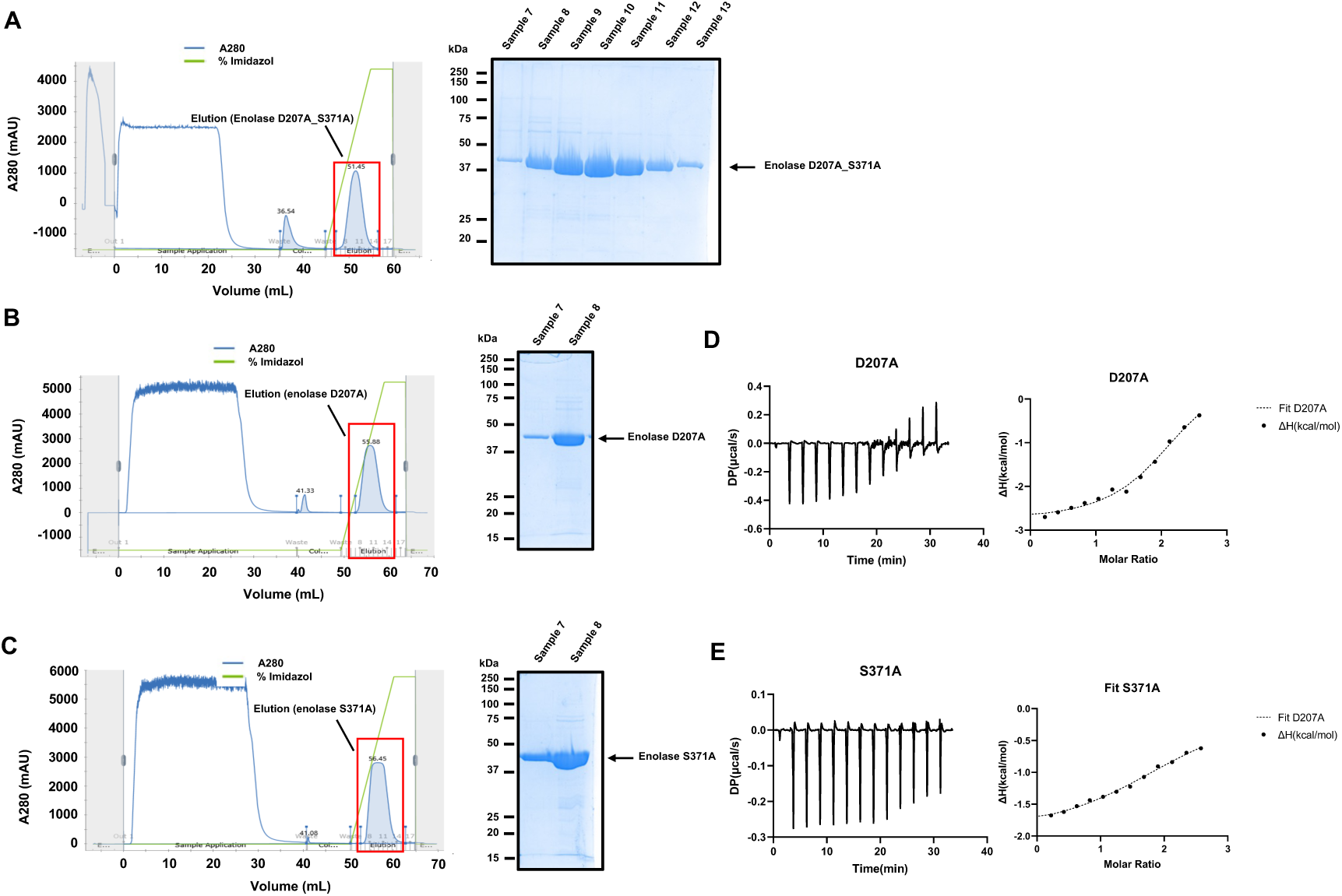
Construction and characterization of *A. baumannii* enolase mutants. (A-. **C)** Purification and SDS-PAGE gel analysis of the purified enolase D207_S371, enolase D207A and enolase S371 using the Histrap FF column. **(E,F)** Isothermal titration calorimetry (ITC) titrations with integrated fitted heat plots of ENOblock binding with enolase (D207A) or enolase (S371A).

**Figure S4.**
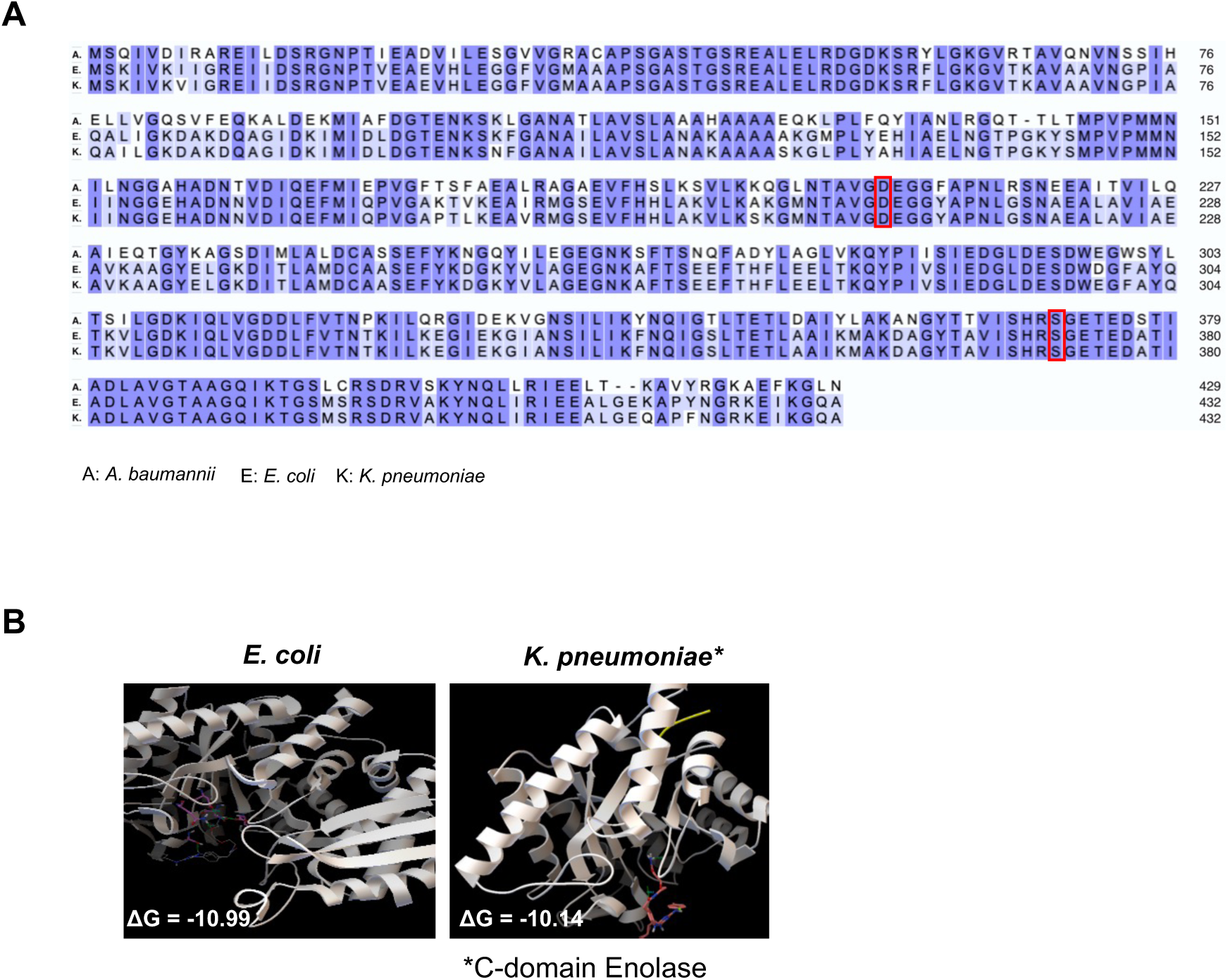
ENOblock acts on *E. coli and K. pneumoniae* through the inhibition of enolase. (**A**) The enolase sequences of *A. baumannii*, *E. coli* and *K. pneumoniae*. (**B**) Structural models generated by docking of ENOblock into *E. coli* and *K. pneumoniae* enolase. ENOblock is displayed as sticks. ΔG: Glide score.

**Figure S5.**
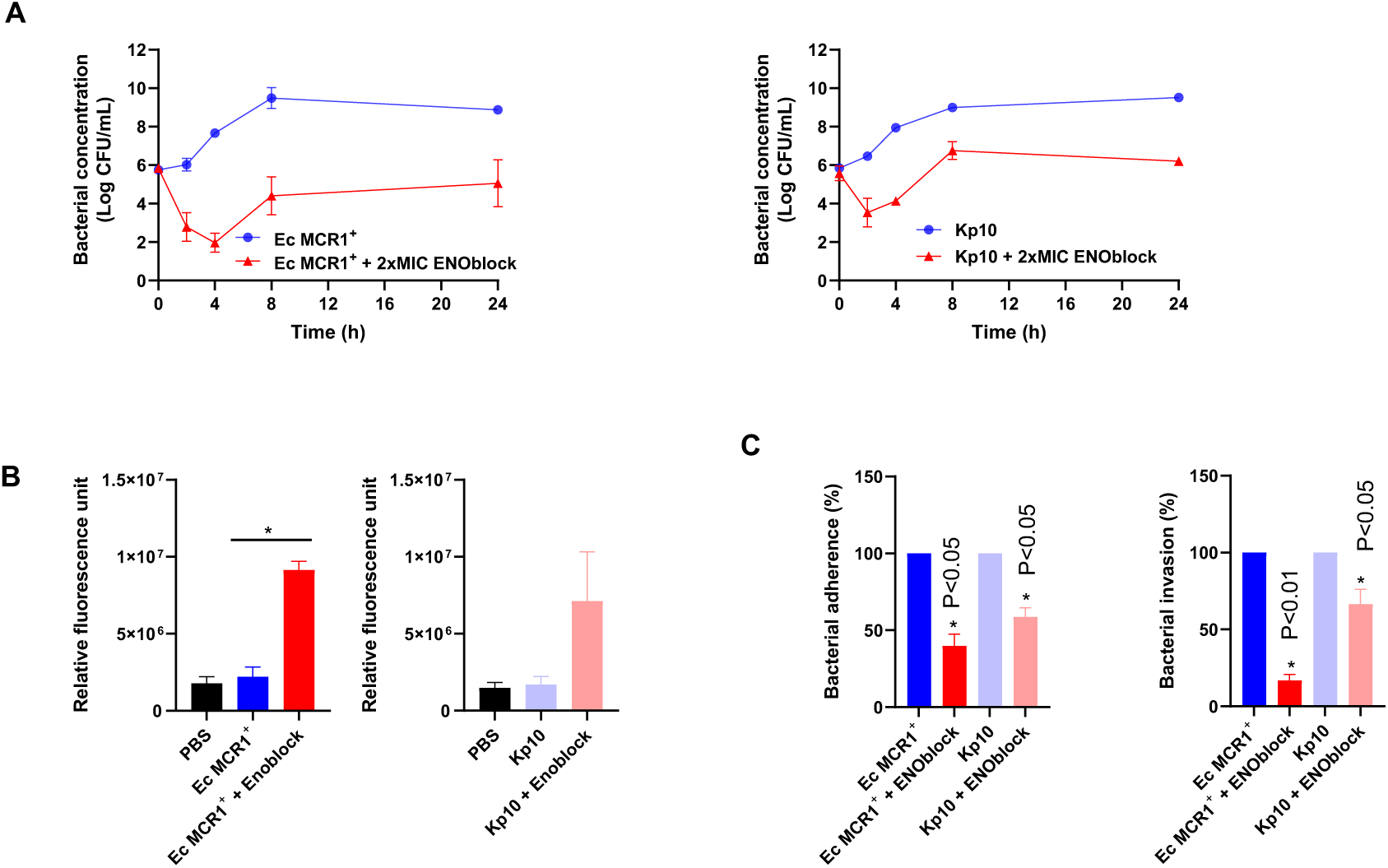
ENOblock is active against *E. coli and K. pneumoniae*. (**A**) Time-kill curves of *E. coli* Ec MCR1^+^ and Kp10 strains in the presence of 2xMIC ENOblock for 24 hours. (B) Membrane permeabilization of *E. coli* Ec MCR1^+^ and Kp10 strains in the presence of 0.5xMIC ENOblock, incubated for 10 min, was quantified by Typhon 334 Scanner. Data are represented as mean ± SEM from three independent replicates and experiments. (C) Analysis of *E. coli* Ec MCR1^+^ and Kp10 strains adhesion/invasion into HeLa cells with (1xMIC) and without ENOblock treatment. The data are presented as means ± SEM, \**P*<0.05: treatment vs no treatment; two-tailed Student’s t-test.

**Figure S6.**
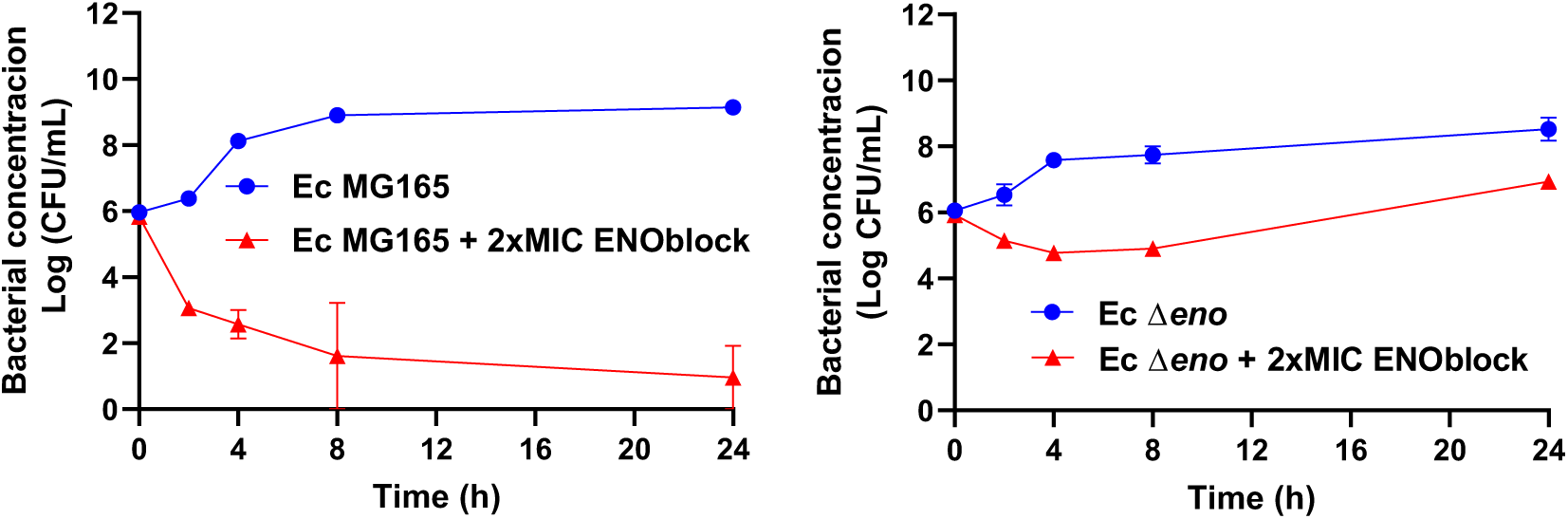
ENOblock acts on *E. coli* through the inhibition of enolase. Time-kill curves of *E. coli* Ec MG165 ^+^ and Ec Δ*eno* strains in the presence of 2xMIC ENOblock for 24 hours

## Notes

### Competing Interest Statement

The authors have declared no competing interest.

## References

1. Morris, S. & Cerceo, E. Trends, epidemiology, and management of multi-drug resistant gram-negative bacterial infections in the hospitalized setting. Antibiotics 9, 196 (2020).

2. Balasubramanian, R. et al. Global incidence in hospital-associated infections resistant to antibiotics: An analysis of point prevalence surveys from 99 countries. PLoS Med. 20, e1004178 (2023).

3. Yelin, I. & Kishony, R. SnapShot: Antibiotic resistance. Cell 172(5), 1136 (2018).

4. Tacconelli, E. et al. Discovery, research, and development of new antibiotics: the WHO priority list of antibiotic-resistant bacteria and tuberculosis. Lancet Infect. Dis. 18, 318–327 (2018).

5. O’Neill, J. Tackling drug-resistant infections globally: final report and recommendations. Review on Antimicrobial Resistance. pp. 1–76 (2016).

6. ECDC and WHO. Antimicrobial resistance surveillance in Europe 2023 - 2021 data. Stockholm: European Centre for Disease Prevention and Control and World Health Organization (2023).

7. Wong, F. et al. Discovery of a structural class of antibiotics with explainable deep learning. Nature 626, 177–185 (2024).

8. Boulaamane, Y., et al. Antibiotic discovery with artificial intelligence for the treatment of Acinetobacter baumannii infections. mSystems 9, e0032524 (2024).

9. Wan, F., Torres, M.D.T., Peng, J. & de la Fuente-Nunez, C. Deep-learning-enabled antibiotic discovery through molecular de-extinction. *Nat*. Biomed. Eng. 8, 854–871 (2024)

10. Miethke, M. et al. Towards the sustainable discovery and development of new antibiotics. Nat. Rev. Chem. 5, 726–749 (2021).

11. Bakker, A.T., et al. Discovery of isoquinoline sulfonamides as allosteric gyrase inhibitors with activity against fluoroquinolone-resistant bacteria. Nat. Chem. In press (2024).

12. Blasco, B. et al. High-throughput screening of small-molecules libraries identified antibacterials against clinically relevant multidrug-resistant *A. baumannii* and *K. pneumoniae*. EBioMedicine 102, 105073. (2024).

13. Zampaloni, C. et al. A novel antibiotic class targeting the lipopolysaccharide transporter. Nature 625, 566–571 (2024).

14. Huang, B. & Zhang, Y. Teaching an old dog new tricks: drug discovery by repositioning natural products and their derivatives. Drug Discov. Today 27, 1936–1944 (2022).

15. Wescott, H.H. et al. Identification of enolase as the target of 2-aminothiazoles in Mycobacterium tuberculosis. Front. Microbiol.9, 2542 (2018).

16. Krucinska, J. et al. Functional and structural basis of E. coli enolase inhibition by SF2312: a mimic of the carbanion intermediate. Sci. Rep. 9, 17106 (2019).

17. Valencia, R., et al. Nosocomial outbreak of infection with pan-drug-resistant Acinetobacter baumannii in a tertiary care university hospital. Infect. Control Hosp. Epidemiol. 30, 257-263 (2009).

18. López-Rojas, R., Jiménez-Mejías, M.E., Lepe, J.A. & Pachón, J. *Acinetobacter baumannii* resistant to colistin alters its antibiotic resistance profile: a case report from Spain. J. Infect. Dis. 204, 1147–1148 (2011).

19. Herrera-Espejo, S. et al. Efficacy of tamoxifen metabolites in combination with colistin and tigecycline in experimental murine models of *Escherichia coli* and *Acinetobacter baumannii*. Antibiotics 13, 386 (2024).

20. Jayol, A., Nordmann, P., Poirel, L. & Dubois, V. Ceftazidime/avibactam alone or in combination with aztreonam against colistin-resistant and carbapenemase-producing Klebsiella pneumoniae. J. Antimicrob. Chemother. 73(**2**), 542–544 (2018).

21. Zhang, J.H., Chung, T.D. & Oldenburg, K.R. A simple statistical parameter for use in evaluation and validation of high throughput screening assays. J. Biomol. Screen. 4, 67–73 (1999).

22. European Committee on Antimicrobial Susceptibility Testing. European antimicrobial breakpoints. Basel: EUCAST, https://www.eucast.org/ast_of_bacteria/mic_determination (2023).

23. Murashko, O.N. & Lin Chao, S. *Escherichia coli* responds to environmental changes using enolasic degradosomes and stabilized DicF sRNA to alter cellular morphology. Proc Natl Acad Sci USA 114, E8025–E8034 (2017).

24. Miró-Canturri, A. et al. Repositioning rafoxanide to treat Gram-negative bacilli infections. J. Antimicrob. Chemother. 75, 1895–1905 (2020).

25. Veljkovic, V. & Slavic, I. Simple general-model pseudopotential. Phys. Rev. Lett. 29, 105–107 (1972).

26. Duran, A., Zamora, I. & Pastor, M. Suitability of GRIND-based principal properties for the description of molecular similarity and ligand-based virtual screening. J. Chem. Inf. Model. 49, 2129–2138 (2009).

27. Veljkovic, V.A. Theoretical Approach to Preselection of Carcinogens and Chemical Carcinogenesis; Gordon & Breach: New York, NY, USA (1980).

28. Schindelin, J., et al. Fiji: an open-source platform for biological-image analysis. Nat. Methods 9, 676–682 (2012).

29. McQuin, C. et al. CellProfiler 3.0: Next-generation image processing for biology. PLOS Biol. 16, e2005970 (2018).

30. Samernate, T. et al. High-Resolution Bacterial Cytological Profiling Reveals Intrapopulation Morphological Variations upon Antibiotic Exposure. Antimicrob. Agents Chemother. 67, e01307–22 (2023).

31. Jiang, C. et al. In silico prediction of chemical neurotoxicity using machine learning. Toxicol. Res. 9, 164–172 (2020).

32. Campello, R.J.G.B., Moulavi, D., Zimek, A. & Sander, J. Hierarchical density estimates for data clustering, visualization, and outlier detection. ACM Trans. Knowl. Discov. Data 10, 1–51 (2015).

33. Wang, Y., Huang, H., Rudin, C. & Shaposhnik, Y. Understanding how dimension reduction tools work: An empirical approach to deciphering t-SNE, UMAP, TriMAP, and PaCMAP for Data Visualization. ArXiv201204456 Cs Stat (2021).

34. Jumper, J. et al. Highly accurate protein structure prediction with AlphaFold. Nature 596, 583–589 (2021).

35. Smani, Y., Dominguez-Herrera, J. & Pachón, J. Association of the outer membrane protein Omp33 with fitness and virulence of *Acinetobacter baumannii*. J. Infect. Dis. 208, 1561–70. 2013

36. Smani, Y. et al. Role of OmpA in the multidrug resistance phenotype of *Acinetobacter baumannii*. Antimicrob. Agents Chemother. 58, 1806–1808 (2014).

37. Parra-Millán, R. et al. Intracellular trafficking and persistence of *Acinetobacter baumannii* requires Transcription Factor EB. mSphere 3, e00106–18 (2018).

38. Díez-Sainz, E. et al. miR482f and miR482c-5p from edible plant-derived foods inhibit the expression of pro-inflammatory genes in human THP-1 macrophages. Front. Nutr. 10, 1287312 (2023).

39. Vila-Farrés, X. et al. Combating virulence of Gram-negative bacilli by OmpA inhibition. Sci. Rep. 7:14683 (2017).

40. Veljkovic, N., Glisic, S., Perovic, V. & Veljkovic, V. The role of long-range intermolecular interactions in discovery of new drugs. Exp. Opin. Drug Disc. 6, 1263–1270 (2011).

41. Sencanski, M. et al. Identification of SARS-CoV-2 Papain-like Protease (PLpro) Inhibitors Using Combined Computational Approach. ChemistryOpen 11(**2**):e202100248 (2022).

42. Smani, Y., McConnell, M.J. & Pachón, J. Role of fibronectin in the adhesion of Acinetobacter baumannii to host cells. PLoS One 7(**4**):e33073 (2012).

43. Martínez-Guitián, M. et al. Antisense inhibition of lpxB gene expression in *Acinetobacter baumannii* by peptide-PNA conjugates and synergy with colistin. J. Antimicrob. Chemother. 75, 51–59 (2020).

44. Nonejuie, P., Burkart, M., Pogliano, K. & Pogliano, J. Bacterial cytological profiling rapidly identifies the cellular pathways targeted by antibacterial molecules. Proc. Natl. Acad. Sci. U.S.A. 110, 16169–16174 (2013).

45. Htoo, H.H. et al. Bacterial cytological profiling as a tool to study mechanisms of action of antibiotics that are active against *Acinetobacter baumannii*. Antimicrob. Agents Chemother. 63, e02310–18 (2019).

46. Htoo, H.H. et al. Mansonone G and its derivatives exhibit membrane permeabilizing activities against bacteria. PloS One 17, e0273614 (2022).

47. Khunsri, I. et al. Roles of qseC mutation in bacterial resistance against anti-lipopolysaccharide factor isoform 3 (ALFPm3). PloS One 18, e0286764 (2023).

48. Lin, L. et al. Azithromycin synergizes with cationic antimicrobial peptides to exert bactericidal and therapeutic activity against highly multidrug-resistant Gram-negative bacterial pathogens. EBioMedicine 2, 690–698 (2015).

49. Ceremuga, I. et al. Enolase-like protein present on the outer membrane of *Pseudomonas aeruginosa* binds plasminogen. Folia Microbiol. 59, 391–397 (2014).

50. Esgleas, M. et al. Isolation and characterization of alpha-enolase, a novel fibronectin-binding protein from *Streptococcus suis*. Microbiology 154, 2668–2679 (2008)

51. Miró-Canturri, A., Ayerbe-Algaba, R., & Smani, Y. Drug repurposing for the treatment of bacterial and fungal infections. Front. Microbiol. 10, 41 (2019).

52. Miró-Canturri, A. et al. Repurposing of the tamoxifen metabolites to treat methicillin-resistant *Staphylococcus epidermidis* and vancomycin-resistant *Enterococcus faecalis* infections. Microbiol. Spectr. 9, e0040321 (2021).

53. Miró-Canturri, A. et al. Repurposing of the tamoxifen metabolites to combat infections by multidrug-resistant gram-negative bacilli. Antibiotics 10, 336 (2021).

54. Miró-Canturri, A. et al. Potential tamoxifen repurposing to combat infections by multidrug-resistant Gram-negative bacilli. Pharmaceuticals 14, 507 (2021).

55. European Committee on Antimicrobial Susceptibility Testing, European antimicrobial breakpoints. Basel: EUCAST (accessed 13 August 2024) (2024).

56. Veljkovic, V. et al. Simple Chemoinformatics criterion using electron donor-acceptor molecular characteristics for selection of antibiotics against multi-drug-resistant bacteria. Discoveries 4(3): e64 (2016).

57. Sullivan, G.J., Delgado, N.N., Maharjan, R., & Cain, A.K. How antibiotics work together: molecular mechanisms behind combination therapy. Curr. Opin. Microbiol. 57, 31–40 (2020).

58. Bojić, T., et al. Virtual screen for repurposing of drugs for candidate influenza a M2 ion-channel inhibitors. Front. Cell. Infect. Microbiol. 9: 67 (2019).

59. López-Rojas, R. et al. Impaired virulence and *in vivo* fitness of colistin-resistant *Acinetobacter baumannii*. J. Infect. Dis. 203, 545–8 (2011).

60. Ayerbe-Algaba, R. et al. The anthelmintic oxyclozanide restores the activity of colistin against colistin-resistant Gram-negative bacilli. Int. J. Antimicrob. Agents 54, 507–512 (2019).

61. Jung, D.W. et al. A unique small molecule inhibitor of enolase clarifies its role in fundamental biological processes. ACS Chem. Biol. 8, 1271–1282 (2013).

62. Canback, B., Andersson, S.G. & Kurland, C.G. The global phylogeny of glycolytic enzymes. Proc. Natl. Acad. Sci. U.S.A. 99, 6097–6102 (2002).

63. Witkowska, D. et al. Antibodies against human muscle enolase recognize a 45-kDa bacterial cell wall outer membrane enolase-like protein. FEMS Immunol. Med. Microbiol. 45, 53–62 (2005).

64. Tamadonfar, K.O. et al. Structure–function correlates of fibrinogen binding by *Acinetobacter* adhesins critical in catheter-associated urinary tract infections. Proc. Natl. Acad. Sci. 120, e2212694120 (2023).

65. Koenigs A, Zipfel PF, Kraiczy P. Translation Elongation Factor Tuf of *Acinetobacter baumannii* is a plasminogen-binding protein. PLoS One 10(**7**), e0134418 (2015).

66. Weng, Y. et al. *Pseudomonas aeruginosa* enolase influences bacterial tolerance to oxidative stresses and virulence. Front. Microbiol. 7, 1999 (2016).

